# The TRIPLE PHD FINGERS proteins are required for SWI/SNF complex-mediated +1 nucleosome positioning and 5’ transcript length determination in Arabidopsis

**DOI:** 10.1101/2022.06.12.495191

**Authors:** Borja Diego-Martin, Jaime Pérez-Alemany, Joan Candela-Ferre, Antonio Corbalán-Acedo, Juan Pereyra, David Alabadí, Yasaman Jami-Alahmadi, James Wohlschlegel, Javier Gallego-Bartolomé

**Author notes:** To whom correspondence should be addressed., Tel: +*34 963877856. “Joint Authors” (Borja Diego-Martin, Jaime Pérez-Alemany).

## Abstract

Eukaryotes have evolved multiple ATP-dependent chromatin remodelers to shape the nucleosome landscape. We recently uncovered an evolutionarily conserved SWItch/Sucrose Non-Fermentable (SWI/SNF) chromatin remodeler complex in plants reminiscent of the mammalian BAF subclass, which specifically incorporates the MINUSCULE (MINU) catalytic subunits and the TRIPLE PHD FINGERS (TPF) signature subunits. Here we report experimental evidence that establishes the functional relevance of TPF proteins for the complex activity. Our results show that depletion of TPF triggers similar pleiotropic phenotypes and molecular defects to those found in *minu* mutants. Moreover, we report the genomic location of MINU2 and TPF proteins as representative members of the plant BAF-like complex and their impact on nucleosome positioning and transcription. These analyses unravel the binding of the complex to thousands of genes where it modulates the position of the +1 nucleosome. These targets tend to produce 5’-shifted transcripts in the *tpf* and *minu* mutants pointing to the participation of the complex in alternative transcriptional start site usage. Interestingly, there is a remarkable correlation between +1 nucleosome shift and 5’ transcript length change suggesting their functional connection. In summary, this study unravels the function of a plant SWI/SNF complex involved in +1 nucleosome positioning and 5’ transcript length determination.

## INTRODUCTION

The genetic material in the eukaryotic nuclei is present in the form of chromatin. The nucleosome is the basic unit of chromatin and it is formed by pairs of histones H3, H4, H2A, and H2B wrapped with approximately 147 base pairs (bp) of DNA (1). This structure represents the first layer of compaction of the genome and has a direct impact on the ability of nuclear proteins to access DNA. Moreover, it serves as a recruitment platform for multiple histone reader proteins (2). Thus, nucleosomes play an important role in the regulation of chromatin affecting diverse processes such as transcription, replication, and repair (3–5). Nucleosomes present a stereotypical distribution relative to the transcription start site (TSS) of genes transcribed by RNA Polymerase II (Pol II) where a strongly positioned nucleosome known as +1 is followed by an array of regularly spaced nucleosomes (+2, +3, etc) and is preceded by a nucleosome depleted region (NDR) that facilitates the access of the transcriptional machinery to the promoter region (6–8). The +1 nucleosome exerts an important function in transcriptional regulation since it is a barrier to Pol II (9,10), promoting Pol II pausing in some species (11–14). Moreover, studies in fungi and animals have shown that the +1 nucleosome can have a negative effect on the ability of the transcriptional machinery to access the TSS and other promoter regulatory regions, leading to changes in gene expression and alternative TSS usage, which in turn can result in altered non-coding transcription and mRNA function (15–21).

Eukaryotes have evolved several ATP-dependent chromatin remodeler complexes to assemble, evict, slide, or restructure nucleosomes (22,23). Among these, the SWItch/Sucrose Non-Fermentable (SWI/SNF) chromatin remodeler family has been extensively studied in diverse eukaryotic model organisms (24–26). These complexes play a major role in the regulation of nucleosome positioning promoting an open NDR (16,27,28) and have a strong impact on cell differentiation processes being frequently mutated in different human cancers (29). The composition of SWI/SNF complexes is diverse and dynamic and is characterized by the presence of one catalytic ATPase subunit and a cohort of scaffold and regulatory subunits (30). There are different evolutionary conserved SWI/SNF subclasses that are characterized by unique signature subunits and that show non-redundant functions (26,29,31). These distinct subclasses are known as SWI/SNF-BAF and RSC-PBAF in fungi and mammals, respectively, and a newer subclass was recently identified in mammals named non-canonical BAF (ncBAF) (32–35).

Plants conserve multiple SWI/SNF subunits and their misregulation can cause strong developmental defects and altered responses to the environment (36,37). The model plant Arabidopsis presents multiple paralogs of these subunits e.g. four ATPases (BRAHMA (BRM), SPLAYED (SYD), MINUSCULE1 (MINU1), and MINU2), four SWI3s (A, B, C, and D), and two SWP73s (A and B), which suggest the formation of multiple variants of the SWI/SNF complex (36). Through a comprehensive evolutionary study of the conservation of SWI/SNF subunits across eukaryotes, we recently reported a model for the evolution of SWI/SNF subclasses that predicts the presence of two major subclasses in plants (35). One is reminiscent of the mammalian BAF-PBAF complexes and incorporates the MINU ATPases while the other is similar to the metazoan ncBAF complex and incorporates the BRM ATPase (35). There is abundant information about the molecular function of subunits of the plant ncBAF-like complex, like BRM (38–41), as well as the bromodomain-containing proteins BRD1,2,13 (BRDs), BRAHMA-INTERACTING PROTEINS (BRIPs), and GRF-INTERACTING FACTORS (GIF) (42–45) signature subunits. However, there is a significant lack of information about the components and molecular function of the plant BAF-like complex. Through proteomics, we recently reported the composition of this complex in Arabidopsis, which incorporates conserved subunits as well as a set of plant-specific subunits of unknown function (35). Among them, we found two paralogs of an evolutionarily-conserved signature subunit –SMARCG-that we named TRIPLE PHD FINGERS (TPF1 and TFP2) and that have not been characterized in plants. These proteins are distant orthologs of the metazoan DOBLE PHD FINGERS(DPF)/PHD FINGER PROTEIN 10 (PHF10) family and the fungal SWP82/Rsc7 proteins (46–48). Functional characterization of the DPF/PHF10 proteins in mammals has demonstrated their important function in the complex through dynamic incorporation upon different cellular contexts and its role in the recruitment of the complex to the chromatin mediated by their tandem PHD histone reader domains (46,49,50). The TPF proteins present three tandem PHD domains as well as a C-terminal Tudor domain, also involved in histone reading, suggesting a possible functional conservation with their animal counterparts (35).

In this study we have explored the function of the plant BAF-like complex through a set of genomic studies that reveal a significant impact of the complex on the +1 nucleosome positioning and the TSS usage. Consistent with a prominent role in the function of the complex, *tpf* mutants present similar pleiotropic phenotypes and molecular defects to those found in mutants of the catalytic subunits of the BAF-like complex.

## MATERIAL AND METHODS

### Plant materials and growth conditions

Plants were grown in a growth chamber with a 16L/8D light cycle (LD) and 22°C. When grown on plates, plants were grown on ½ MS medium with vitamins (Duchefa) 0.8% agar pH 5.7 in a 16L/8D light cycle and 22°C. Transgenic lines were selected on MS medium supplemented with BASTA (6,67 μg/mL) or Hygromycin (35 μg/mL). For the root growth experiment, seeds were stratified for 7 days (darkness 4°C) and plants were grown on MS vertical plates for 12 days under LD photoperiod. The T-DNA insertion lines used in this study were the following: *minu1-2* (GK-146E09), *minu2-1* (SALK_057856C), *tpf1-1* (SALK_010411C), *tpf1-2* (WiscDsLox385F06), *tpf2-1* (SALK_141512C), and *tpf2-2* (SAIL_201_D01).

### Transgenic lines

A genomic fragment of TPF2, including promoter until the codon before the stop, was amplified from genomic DNA and cloned into a pENTR/D plasmid by InFusion (Takara) to generate pENTR-gTPF2. The TPF2 fragment was transferred by LR reaction into a modified pEarleyGate302 that contains a 3xFLAG tag downstream of the gateway cassette to generate pEG-gTPF2-3xFLAG. This construct was transformed into *Agrobacterium tumefaciens* strain *GV3101 C58C1,* which was used to transform the *tpf2-1* mutant by the floral dip method. A genomic fragment of MINU2, including promoter until the codon before the stop, and with an inserted XhoI site right upstream of the start codon was amplified from genomic DNA in two separate PCR reactions. These fragments were combined to amplify the full genomic sequence that was cloned into a pENTR/D plasmid by InFusion (Takara) to generate pENTR-gMINU2. A 3xFLAG tag was amplified and cloned into the unique XhoI site of pENTR-gMINU2. The 3xFLAG-gMINU2 fragment was transferred by LR reaction into a pMDC123 (51) to generate pMDC123-3xFLAG-gMINU2. This construct was transformed into *Agrobacterium tumefaciens* strain *GV3101 C58C1,* which was used to transform the *minu2-1* mutant by the floral dip method.

### Root meristem microscopy

Roots from plants grown 12 days on vertical plates were incubated with 10 μg/mL Propidium Iodide and visualized in a confocal microscope Axio Observer 780 (Zeiss).

### Yeast two-hybrid (Y2H)

Gateway-compatible vectors that incorporate the SWI/SNF subunits were either requested from ABRC as bacterial stabs (SHH2 (U17329), SWI3A (U16949), SWI3B (G20908), SWP73B (G15375), LFR (G20071), BSH (G82285), ARP4 (G23361), ARP7 (U24661), PSA2 (U25108), BDH1 (G50477), OPF2 (G68780), TPF2 (TOPO-U19-C04), MINU2 (TOPO-U19-F11)) or cloned from cDNA into a pENTR/D or pDONR207 plasmid (PSA1 and BDH2, see Supplementary Data 1 for primer information). The pENTR-TPF1, pENTR-MINU1, and pENTR-BRD5 plasmids, containing their respective full-length CDS sequences, were synthesised from Genescript. The pENTR-TPF1 was mutated to generate the pENTR-N-TPF1 (TPF1 amino acids 1-422). The C-TPF1 fragment (TPF1 amino acids 423-697) was amplified from pENTR-TPF1 and cloned into a pDONR207 plasmid by PB reaction (Invitrogen). Plasmids were transferred to pGADT7 and pGBKT7 vectors (Clontech) by LR reaction (Invitrogen).

Yeasts were grown in SD media containing 6.7 g/L DifcoTM Yeast Nitrogen Base w/o amino acids (Becton Dickinson S.A.), and 1.4 g/L Yeast Synthetic Drop Out Medium Supplements (Sigma) pH 5.8, supplemented with 5% glucose, 10 μg/mL (Ura, Trp, His) and 50 μg/mL Leu. pGADT7 and pGBKT7 plasmids (Clontech) were transformed, following the lithium acetate/single-stranded carrier DNA/polyethylene glycol method, into the haploid strains *Y187* and *Y2HGold,* which were selected in SD media devoided of Leu and Trp, respectively. Diploid yeasts were obtained by mating, which was carried out overnight in YPD supplemented with 2% glucose, and the generated diploid cells were selected in SD/-Leu-Trp plates. Protein interaction was tested based on the complementation of the histidine autotrophy, by dropping serial dilutions into SD/-Leu-Trp-His plates.

### Immunoprecipitation mass spectrometry (IP-MS)

Immunoprecipitation followed by mass spectrometry was done as previously described (35). Briefly, 8 g of inflorescences from one untransformed Col-0 replicate, two independent lines of TPF2-FLAG and two independent lines of MINU2-FLAG, were ground in liquid nitrogen and resuspended in 40 ml IP buffer (50 mM Tris pH 7.6, 150 mM NaCl, 5 mM MgCl_2_, 10% glycerol, 0.1% NP40, 0.5 mM DTT, 1 mM PMSF, 1 μg/μL pepstatin, and 1× Complete EDTA-Free (Sigma). Samples were filtered with one layer of Miracloth (Merck, cat#475855), and homogenized with a douncer (10 times soft, 10 times hard). Then, centrifuged at 4°C for 10 min at 10,000 x *g* and filtered again using a 40 μm cell strainer. 200 μl of Anti-FLAG M2 magnetic beads (Sigma, cat#M8823), previously blocked with 5% BSA, were added to the samples followed by 3 hours rotating at 4°C. The samples were washed 4 times with IP buffer and two times with IP buffer without NP40. Samples were eluted with 300 μl of 250 μg/ml 3xFLAG peptide (Sigma, cat#F4799) in IP buffer without NP40 for 30 min at 25°C. This step was repeated one more time incubating samples for 15 min at 37°C. TCA was added to a final concentration of 20% and incubated for 30 min on ice followed by a 30 min 4°C centrifugation at 12,000 x *g*. Subsequently, samples were washed three times with 250 μl of cold acetone and the pellet was air-dried.

For the purpose of proteomics, proteins were reduced and alkylated using 5 mM Tris (2-carboxyethyl) phosphine and 10 mM iodoacetamide, respectively. Protein digestion was achieved by sequential addition of endopeptidase Lys-C (BioLabs) and trypsin (Pierce™) at 1:100 enzyme/protein ratio was done to digest proteins followed by an incubation at 37°C overnight. Formic acid to 5% (v./v.) final concentration was added to quench the samples. Finally, C18 pipette tips (Thermo Scientific, cat# 87784) were used for desalting prior to LS-MS/MS analysis and reconstituted in 5% formic acid before being analyzed by LC-MS/MS. A 25 cm long, 75 μm ID fused-silica capillary that was packed in-house with bulk ReproSil-Pur 120 C18-AQ particles as described elsewhere (52) was used to fractionate online the peptide mixtures. Peptides were subjected to a 140 min water-acetonitrile linear gradient in 6-28% buffer B (acetonitrile solution with 3% DMSO and 0.1% formic acid) at a flow rate of 200 nl.min^-1^, which was further increased to 35% followed by a rapid ramp-up to 85% using a Dionex Ultimate 3000 UHPLC (Thermo Fisher Scientific). The eluted peptides were then ionized via nanoelectrospray ionization. An *Orbitrap Fusion™* Lumos™ *Tribrid™* Mass Spectrometer (Thermo Fisher Scientific) was used to acquire the mass spectrometry data with an MS1 resolution of 120,000 followed by sequential MS2 scans at a resolution of 15,000. Raw data were searched against the TAIR Arabidopsis reference proteome. Default settings for LFQ analysis using MaxQuant 1.6.17.0 software were applied to calculate the Label-free quantitation (LFQ) intensities as described previously by (53). The *LFQ* intensities were log2-transformed and used for further processing.

Shown interactors in Table 1 followed the criteria LFQ control < LFQ transgenic/10 in all TFP2-3xFLAG and 3xFLAG-MINU2 replicates. Values for TPF1 were included manually since it was absent in the TPF2-3xFLAG experiments. The TPF1 and SHH2 results were previously published (35) and are shown to compare with the new datasets.

**Table 1.** Plant BAF-like complex subunits identified through IP-MS experiments using TPF2, MINU2, TPF1 and SHH2 as baits.

| Name | Protein ID |  | TPF2-3xFLAG |  | 3xFLAG-MINU2 |  | TPF1 experiment 1 |  |  | SHH2 experiment 1 |  |  |
| --- | --- | --- | --- | --- | --- | --- | --- | --- | --- | --- | --- | --- |
|  |  | Col-0 | #9 | #15 | #7 | #10 | Col-0 | #11 | #4 | Col-0 | #10 | #21 |
| TPF2 | AT3G08020 | - | 38 | 36 | 33 | 33 | - | - | - | - | 28 | 27 |
| SWI3B | AT2G33610 | - | 34 | 33 | 33 | 33 | - | 29 | 26 | - | 30 | 29 |
| SWI3A | AT2G47620 | - | 33 | 32 | 32 | 33 | - | 28 | 26 | - | 30 | 30 |
| MINU1 | AT3G06010 | - | 33 | 32 | 30 | 31 | - | 28 | 26 | - | 30 | 29 |
| ARP4 | AT1G18450 | 25 | 33 | 32 | 32 | 33 | 25 | 30 | 31 | 27 | 30 | 29 |
| PSA1 | AT1G32730 | - | 33 | 32 | 32 | 34 | - | 29 | 28 | - | 29 | 28 |
| ARP7 | AT3G60830 | - | 33 | 32 | 32 | 34 | 24 | 32 | 32 | 28 | 31 | 31 |
| CHC1 | AT5G14170 | - | 33 | 32 | 32 | 33 | - | 28 | 26 | 25 | 30 | 29 |
| LFR | AT3G22990 | - | 33 | 32 | 32 | 33 | - | 25 | 23 | 23 | 30 | 28 |
| PSA2 | AT1G06500 | - | 32 | 31 | 31 | 32 | - | 30 | 30 | - | 28 | 28 |
| BDH2 | AT5G55210 | - | 32 | 31 | 32 | 31 | - | - | - | - | 27 | - |
| OPF1 | AT1G50620 | - | 32 | 31 | 31 | 32 | - | 29 | 28 | - | 33 | 33 |
| BRD5 | AT1G58025 | - | 32 | 31 | 31 | 32 | - | 27 | 25 | - | 29 | 27 |
| MINU2 | AT5G19310 | - | 31 | 31 | 33 | 34 | - | - | - | - | 27 | 24 |
| BSH | AT3G17590 | - | 32 | 30 | 31 | 32 | - | 25 | - | - | 30 | 29 |
| BDH1 | AT4G22320 | - | 31 | 30 | 31 | 32 | - | 29 | 30 | - | 27 | 27 |
| SHH2 | AT3G18380 | - | 30 | 29 | 29 | 31 | - | 29 | 29 | - | 35 | 36 |
| OPF2 | AT3G20280 | - | 30 | 28 | 30 | 31 | - | - | - | - | 30 | 30 |
| TPF1 | AT3G52100 | - | - | - | 34 | 35 | - | 36 | 36 | - | 31 | 30 |

Data represents Log2LFQ intensity values of the plant BAF-like complex subunits identified by IP-MS experiments using two independent lines of TPF2-3xFLAG (9 and 15) and 3xFLAG-MINU2 (7 and 10) compared to untransformed Col-0 controls. Shown interactors followed the criteria LFQ control < LFQ transgenic/10 in both transgenic lines compared to Col-0 control and were selected based on the previous identification of plant BAF-like subunits using TPF1 and SHH2 as baits (35). Values from representative TPF1 and SHH2 IP-MS experiments are shown (35). Raw data for the TPF2 and MINU2 IP-MS experiments can be found in Supplementary data 12.

### RNA extraction, qRT-PCR and RNA-seq libraries

Total RNA from inflorescences was extracted with the NucleoSpin RNA Plant kit (Macherey-Nagel) following the manusfacturer’s protocol and adding an on-column DNase treatment. For the RNA-seq experiments, RNA was extracted from inflorescences of Col-0, *minu1-2 minu2-1,* and *tpf1-1 tpf2-1* plants (3 biological replicates per background) and sent to BGI to prepare strand-specific mRNA libraries that were sequenced by DNBSEQ high-throughput platform as PE100 reads. For the characterization of alternative TSS usage, 1 μg of the same RNA as the one used for RNA-seq experiments was used for cDNA synthesis using the Superscript IV kit (Invitrogen) and a modified oligodT (JG541) described in (54), followed by RNAse H treatment according to the manufacturer’s protocols. For the *TPF* expression in the T-DNA lines, 1 μg of total RNA extracted from 13-day-old seedlings was used for cDNA synthesis using the NZY First-Strand cDNA Synthesis kit (NZYTech) following the manufacturer’s protocol. Primer information can be found in Supplementary Data 1. The fold change was calculated against a housekeeping gene (*PP2A*) following the AACt method (55).

### Chromatin Immunoprecipitation (ChIP-seq)

The chromatin immunoprecipitation protocol was done as previously described with minor changes (56). Briefly, 1 g of inflorescences were ground in liquid nitrogen and fixed in Nuclei Isolation buffer containing 1% formaldehyde for 10 min at RT. In the case of MINU2-FLAG ChIPs, samples were first fixed in Nuclei Isolation buffer containing 1.5 mM EGS for 20 minutes at RT followed by the addition of 1% formaldehyde and incubation for 10 additional minutes. Reactions were stopped with glycine followed by nuclei isolation and chromatin shearing using a Bioruptoptor pico (Diagenode). Chromatin was immunoprecipitated overnight at 4°C with the following antibodies: Anti-FLAG M2 (5μl/ChIP used, F1804, Sigma), H3K4me3 (5μl/ChIP used, 04-745, Millipore), H3K36me3 (10 μl/ChIP used, Ab9050, Abcam), H3K27me3 (10 μl/ChIP used, 07-449, Millipore), H3 (5 μl/ChIP used, Ab1791, Abcam), H2A (10 μl/ChIP used, Ab13923, Abcam), H2A.Z (3 μl /ChIP used), and PanH3Ac (5 μl/ChIP used, 39140, Active motif). Complexes were captured with a 1:1 mixture of magnetic Protein A and Protein G Dynabeads (Invitrogen) for 3 h at 4°C, washed with low salt, high salt, LiCl, and TE buffers for 10 min each at 4°C, and eluted for 2 × 20 min at 65°C with elution buffer. Reverse crosslink was done overnight at 65°C, followed by proteinase K treatment at 45°C for 5 h. DNA was purified using Phenol:Chloroform:Isoamyl Alcohol 25:24:1 (Fisher Scientific) and precipitated with GlycoBlue (Invitrogen) and NaAc/EtOH overnight at −20°C. DNA was resuspended in 75 μl of elution buffer. Libraries for sequencing were prepared using the Ovation Ultra Low System V2 1-16 kit (NuGEN) following the manufacturer’s instructions. Libraries were sequenced in a HiSeq 4000 (Histone and TPF1 experiments) and HiSeq 2500 (TPF2 and MINU2 experiments) as SE50 reads.

### Micrococcal Nuclease assay (MNase-seq)

0.35 g of inflorescences from Col-0, *minu1-2 minu2-1,* and *tpf1-1 tpf2-1* (2 biological replicates per background) were ground and resuspended in 20 ml of Isolation Buffer 1 (20 mM Tris pH8, 0.3 M sucrose, 5 mM MgCl2, 0.2% Triton X-100, 5 mM BME, 35% glycerol, 0.1 μM PMSF, Complete mini -EDTA). After filtering through 1 layer of Miracloth (Merck, cat#475855), samples were spun at 3200 rpm 4°C for 20 min. The pellet was resuspended in 1 ml of HBB buffer (25 mM Tris–HCl, pH 7.6, 0.44 M sucrose, 10 mM MgCl_2_, 0.1% Triton X-100 and 10 mM β-mercaptoethanol, 0.1 μM PMSF, Complete mini –EDTA) and spun at 3200 rpm at 4°C for 10 min. Then, the pellet was resuspended in 1 ml of MNB buffer (10% sucrose, 50 mM Tris-HCl, pH 7.5, 4 mM MgCl_2_, and 1 mM CaCl_2_) and spun at 3200 rpm 4°C for 10 min. Finally, the pellet was resuspended in 900 μl of MNB buffer and 180 μl aliquots were digested with different amounts of Micrococcal Nuclease (N3755, Sigma) (0, 1, 3, 5, and 8U) followed 10 min incubation at 37°C, quickly mixing samples every 3 min. Reactions were stopped by the addition of 20 μL of 0.5 mM EDTA. 4 μl of DNAse-Free RNAse (10 mg/ml) were added followed by incubation at 37°C for 1 h. 1 μl proteinase K (20 mg/ml) was added followed by incubation at 45°C for 1 h. DNA was extracted with Phenol:Chloroform:isoamyl followed by chloroform extraction and precipitation with NaOAc/glycogen/ethanol at −20°C overnight. Samples were centrifuged at 13,000 × *g* for 15 min at 4°C, washed with 70% ethanol and resuspended in 30 μl elution buffer. Samples were run in a 2% agarose gel to recover mononucleosomal DNA (approx. 150 bp) from digestions that presented 20% di-nucleosome and 80% of mono-nucleosome. Gel bands were cut and purified with Zymo gel extraction kit. Libraries for sequencing were prepared at BGI following the BGISEQ-500 ChIP-Seq library preparation protocol and were sequenced by DNBSEQ high-throughput platform as PE100 reads.

### ChIP-seq data analysis

Read quality was checked with FastQC v0.11.9. Reads were then mapped to the TAIR10 genome using Bowtie2 v2.4.4 (57) with default parameters. Duplicates were marked with picard MarkDuplicates v2.26.6 (http://picard.sourceforge.net/). Reads that were either unmapped, duplicated or multimapped were discarded using the samtools (v1.14) (58) view command with options “-F 4 -F 1024 -q 5”. Chipseq-greylist v1.0.2 (https://github.com/roryk/chipseq-greylist) was used to detect anomalous enrichment in control samples. These regions were removed from a TAIR10 chromosome bedfile with bedtools v2.30.9 (59) complement to build a bedfile of included regions. Reads that overlapped the included regions were counted to calculate Reads Per Million (RPM) normalization factors using samtools v.1.14. Fragment sizes of ChIP libraries were estimated with phantompeakqualtools v1.2.2. (60) ChIP coverage bedgraphs were built using the bedtools (v2.30.9) genomecov command, specifying “-scale <RPM factor> -fs <fragment size>“ as parameters. These files were then converted to bigwig using bedGraphToBigWig from UCSC tools. ChIP bigwigs were log2 ratio-normalized against their controls with the bigwigCompare command from deepTools v3.5.1. (61). Coverage and log2 ratio bigwigs from replicates were averaged using wiggletools (v1.2.11) mean (62). Histone PTM ChIP-seqs were compared to a previously published dataset (63) to support the reproducibility of our data.

Peaks were called using MACS v2.7.1 (64) with options “--keep-dup all -g 119482427 --nomodel --extsize <fragment size> --call-summits”. Intersections between peaks were performed using the merge command from bedtools v2.30.9 (59). Reads overlapping common peaks were counted using the multiBamSummary command from deeptools v3.5.1. (59). Venn Diagrams were drawn using eulerr v6.1.1 and the correlation between replicates was calculated using the cor function in R v4.1.2. Peak summits were annotated with genomic features and genes by intersecting them against the Araport11 annotation with the intersect command from bedtools v2.30.9 (59). Genes that overlapped summits or were less than 200 bp away from the TSS were considered as targets. Conversely, genes that did not meet these criteria or overlapped enriched regions in ChIP controls were categorized as not targets.

### MNase-seq data analysis

Read quality control and alignment were performed as described for ChIP-seq data analyses. Fragment size distributions were obtained with the samtools stats command from samtools v1.14. As in ChIP-seq analyses, regions with anomalous enrichment were detected with chipseq-greylist v1.0.2 and removed from a TAIR10 chromosome bedfile using bedtools (v2.30.9) (59) complement. The resulting genomic intervals were converted to wig using a python script. DANPOS v2.2.2. (65) was used to obtain normalized coverage wigs and poisson difference wigs, as well as to carry out analyses of differential nucleosome dynamics. The dpos.py command of DANPOS was run with parameters “--save 1 --paired 1 --span 1 --clonalcut 1e-10 --nor_regions_file <included regions>”. Coverage wig files were converted to bedgraph format using the write_bg command from wiggletools v1.2.11 (62), and these were converted to bigwig with the bedGraphToBigWig command from UCSC tools. Additional MNase-seq raw files from (66) were downloaded and processed to evaluate the similarity of our dataset to previously published data.

Nucleosome bedfiles were built from the output excel files of DANPOS using python scripts. Dyad summits were annotated to their closest gene using the bedtools v2.30.9 (59) closest command and the Araport11 annotation. The +1 nucleosome was identified as the first nucleosome with significant occupancy (summit p-value < 0.05) downstream of the TSS position. To confidently detect differences in nucleosome occupancy and fuzziness, only nucleosomes with significant occupancy in at least one condition were considered. Furthermore, to evaluate position shifts, we only considered those that displayed significant occupancy (summit p-value <0.05) and fuzziness (fuzziness p-value <0.05) in all conditions. This is because nucleosome shift values between not well-positioned nucleosomes tend to be noisy.

### RNA-seq data analysis

Quality control of reads was first performed with FastQC v0.11.9. Reads were then aligned to the TAIR10 genome with STAR v2.7.9a (67). Gene counts were obtained with featureCounts v2.0.1 (68) using the Araport11 annotation. Differential expression analyses were carried out with DESeq2 v1.32.0. The PCA was computed with the plotPCA function over r-log normalized counts from DESeq2 (69). The lfcShrink function was used to obtain differential expression scores with default parameters. Genes with adjusted p-values < 0.05 and |log2FoldChange| > log2(1.5) were considered as differentially expressed. Gene Ontology enrichment analyses were performed over these genes using clusterProfiler v4.0.5 (70) and org.At.tair.db v3.13.0. Coverage bedgraph files were built using bedtools v2.30.9 (59) genomecov with parameters “-bga -split -scale <size factor>”, in which size factors were normalization factors estimated by DESeq2. These were converted to bigwigs using the bedGraphToBigWig command from UCSC tools. The wiggletools v1.2.2 (62) mean command was used to average bigwigs of biological replicates. Finally, log2 ratio bigwigs were built using bigwigCompare from deepTools v3.5.1 (61).

### Data visualization

Statistical graphics such as bar plots, scatter plots, and boxplots were drawn using ggplot2 v3.3.6. Coverage bigwigs were visualized in the IGV browser and screenshots were rendered using SparK v2.6.2. Heatmaps and metaplots were obtained using the computeMatrix, plotHeatmap, and plotProfile commands from deeptools v3.5.1. TSS reference positions for metaplots and screenshots were obtained from an analysis performed in (71) (https://github.com/Maxim-Ivanov/Kindgren_et_al_2019/blob/master/Adjusted_Araport11/genes_araport_adj.bed.gz) which relies on TSS-seq and DR-seq experiments to precisely annotate TSSs.

## RESULTS

### *tpf* mutants have a strong impact on plant development and reproduction

The model plant Arabidopsis possesses two paralogs of the TPF proteins named TPF1 and TPF2 which were previously shown to belong to a plant BAF-like complex in Arabidopsis (35). To study the impact of TPF proteins on plant development, we identified T-DNA insertion lines for the *TPF1* and *TPF2* genes. For *TPF1,* we identified two lines with insertions located in the first exon *(tpf1-1)* and the first intron *(tpf1-2)* (Supplementary Figure 1A). For *TPF2,* we identified two lines, *tpf2-1* and *tpf2-2,* where the T-DNA was inserted in the 11th exon and the intron between exons 10 and 11, respectively (Supplementary Figure 1A). The expression of *TPF1* in the *tpf1* mutant lines was reduced compared to WT (Supplementary Figure 1B, Supplementary Data 2). In the case of *TPF2*, the *tpf2-1* allele caused reduced expression while *tpf2-2* did not show reduced levels compared to wild-type Col-0 plants (WT). According to a public database (www.genevestigator.com) the overall expression pattern of *TPF* genes is similar, showing medium-high expression levels and a stronger difference in senescent tissue (Supplementary Figure 1C). Single mutants of *TPF1* and *TPF2* did not show any apparent developmental differences compared to WT plants (Supplementary Figure 2A). However, combinations of the *TPF1* and *TPF2* alleles resulted in two different mutant phenotypes (Supplementary Figure 2B,C). While combinations of *tpf1-1* and *tpf1-2* with *tpf2-1* resulted in dwarf plants with strong developmental phenotypes, double mutants incorporating the *tpf2-2* allele showed milder defects, suggesting that *tpf2-2* is a weaker allele (Supplementary Figure 2B,C). Thus, further phenotypical characterizations were performed in the strong combination *tpf1-1 tpf2-1* and the weak combination *tpf1-2 tpf2-2,* hereafter named strong and weak *tpf1/2* mutants, respectively. Seedlings of the strong and weak *tpf1/2* mutants were smaller and developmentally delayed and presented shorter roots compared to WT plants (Figure 1A,B), which is probably the consequence of shorter meristems with fewer numbers of cells (Supplementary Figure 3A, Supplementary Data 3). Interestingly, the niche of meristematic cells above the quiescent center incorporated high amounts of propidium iodide (PI) in the strong *tpf1/2* mutant (Figure 1C, Supplementary Figure 3B). This phenotype normally occurs in damaged or dead cells (72) thus suggesting that this pool of cells might have increased DNA damage response in the *tpf1/2* mutant. Adult plants of the strong and weak combinations were smaller and developmentally delayed (Figure 1D,E). The strong *tpf1/2* mutant showed pleiotropic defects with variable penetrance on flower development. Among the most frequent phenotypes were flowers with extra petals, open carpels, protuberances striking out of the stigma (Figure 1F), and defective anther dehiscence (Supplementary Figure 3C). Moreover, it produced small siliques with no seeds (Figure 1G), although very few seeds could be recovered in some mutant plants. The siliques from *tpf1-1 (+/-) tpf2-1 (-/-)* plants presented a high number of abortions and the segregation of double mutant plants was smaller than expected (3:0.38 +/- 0.07 observed; 3:1 expected; Average and standard error calculated from three independent *tpf1-1 (+/-) tpf2-1 (-/-)* populations) (Figure 1H, Supplementary Figure 3D, Supplementary Data 4). In order to gain further insight into the sterility phenotype of the strong *tpf1/2* mutant, we performed reciprocal crosses with the WT (Supplementary Figure 3E, Supplementary Data 5). When WT pollen was used to pollinate mutant pistils, siliques did not elongate and just 4 seeds were recovered out of 28 crosses (Supplementary Figure 3E). On the contrary, when mutant pollen was used on WT pistils, siliques elongated and produced viable seeds (Supplementary Figure 3E), suggesting that the fertilization problem is mainly associated to the female gametophyte. This sterility phenotype is further enhanced by the dehiscence defects in the mutant anthers at the moment of anthesis when self-pollination takes place (Supplementary Figure 3C). In summary, this data shows that the TPF proteins are functionally redundant in the control of several aspects related to plant development and reproduction. Importantly, these phenotypes are reminiscent of the ones reported for double mutants of the functionally redundant MINU1 and MINU2 proteins (73) (Supplementary Figure 2D and 4). This is consistent with the idea that TPF proteins are important players in the overall function of the plant BAF-like complex.

**Figure 1.**
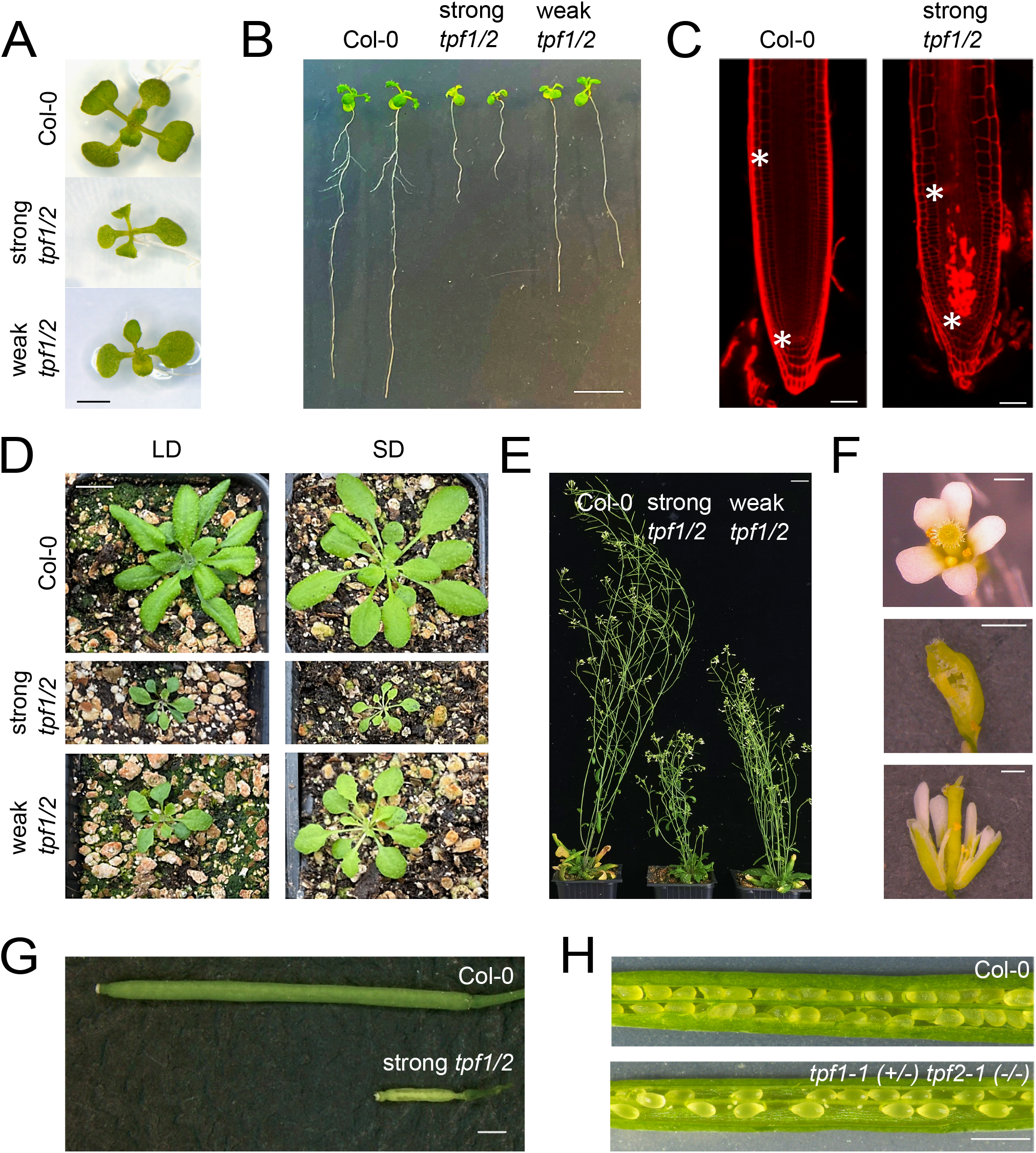
*tpf* mutants have a strong impact on plant development and reproduction. A-B) *In vitro*-grown twelve-day-old seedlings of strong and weak *tpf1/2* mutants show reduced aerial and radicular growth, scale bars: 2 mm and 1 cm, respectively. C) Confocal images of twelve-day-old root tips of Col-0 and the strong *tpf1/2* mutant stained with propidium iodide (PI). Asterisks indicate meristem length. A strong PI signal accumulates above the quiescent center in the *tpf1/2* mutant. Defects in the division plane can be observed in the strong *tpf1/2* mutant. scale bar: 50 μm. D) Top view of three-week-old Col-0, strong, and weak *tpf1/2* mutants grown on soil in long-day conditions and four-week-old plants of the same backgrounds grown on short-day conditions, scale bar: 1 cm. E) Seven-week-old Col-0, strong, and weak *tpf1/2* mutants grown on soil in long-day conditions, scale bar: 2 cm. F) Representative images of the developmental defects found in flowers of the strong *tpf1/2* mutant. Extra petals (upper), open carpels (middle), and protuberances in the stigma (lower), scale bar: 0.5 mm. G) Representative image of siliques of Col-0 and strong *tpf1/2* mutant plants, scale bar: 1 mm. H) Representative images of open siliques of Col-0 and *tpf1-1 (+/-) tpf2-1 (-/-)* showing ovule abortion, scale bar: 1 mm.

### TPF proteins are members of a BAF-like complex in plants

Through IP-MS experiments, we recently showed the composition of a BAF-like SWI/SNF complex in plants (35). To further confirm the complex composition and the mutually exclusive presence of TPF paralogs, we performed additional IP-MS experiments in inflorescences of Arabidopsis using transgenic lines expressing 3xFLAG-tagged TPF2 and MINU2 proteins. Importantly, these transgenes were able to complement the strong *tpf1/2* and the *minu1-2 minu2-1 (minu1/2)* double mutant backgrounds, respectively (Supplementary Figure 4). These experiments confirmed the presence of a well-defined BAF-like complex that incorporates one TPF subunit and specifically the MINU1 and MINU2 ATPases, as well as a set of known and several uncharacterized subunits (Table 1). The mammalian TPF distant ortholog DPF2 directly interacts with several subunits of the SWI/SNF complex including ARID1 (plant LFR) and SMARCD (plant SWP73) (31). We tested the direct interaction between TPF1 and all the identified subunits in the complex by Y2H. For this experiment we split TPF1 into an N-terminal (N-TPF1) fragment that includes the 3xPHD domains and a C-terminal fragment that includes the Tudor domain (C-TPF1) (Supplementary Figure 5A). The N-TPF1 fragment was able to interact with SWP73B and LFR (Supplementary Figure 5B), suggesting conservation of the TPF1 position within the complex compared to its mammalian counterpart. Surprisingly, the same experiment using the TPF1 full-length and C-TPF1 fragments did not result in any positive interactions, probably due to steric impediments or misfolding of the fusion proteins.

### TPF and MINU2 proteins locate over the 5’regions of thousands of genes

The genome-wide location of the BAF-like complex has not been reported in plants. We profiled the location of the MINU2 ATPase, as a core member of the complex, as well as the location of the TPF1 and TPF2 proteins to study whether chromatin-bound BAF-like complex incorporates TPF proteins at all targets. For this purpose, we performed ChIP-seq experiments with two independent transgenic lines expressing 3xFLAG-tagged TPF1, TPF2, and MINU2 (Supplementary Data 6). The two lines tested showed a highly reproducible profile (Supplementary Figure 6A,B). We found 14419 peaks for TPF1, 15337 peaks for TPF2, and 12169 peaks for MINU2 (Figure 2A). Consistent with their redundant function at the phenotypical level, the overlap of TPF peaks was very high (Figure 2A,B,C). These peaks were mostly located over the region downstream of the TSS of thousands of genes (Figure 2B,C,D, Supplementary Data 7). The overlap between MINU2 and TPF peaks was also very high, although the position of MINU2 position was slightly shifted towards the promoter region, perhaps reflecting the different positions occupied by TPF and the ATPase within the complex (Figure 2A,B,C,D). The target genes were enriched with epigenetic marks related to transcription activation such as H3K4me3, H3 pan acetylation, H3K36me3, and H2A.Z (over the TSS regions) while non-target genes were enriched in silencing marks such as H3K27me3 and H2A.Z (over gene body regions) (Figure 2E, Supplementary Figure 7A,B, Supplementary Data 6). Consistently, direct targets were more expressed than non-targets (Figure 2F) and the accumulation of the complex positively correlated with gene expression (Figure 2G). In summary, the plant BAF-like complex locates over the 5’ of euchromatic expressed genes with an epigenome favourable for transcription. Moreover, TPF1 and TPF2 share a large number of targets with MINU2 suggesting they are core members of the chromatin-bound BAF-like complex.

**Figure 2.**
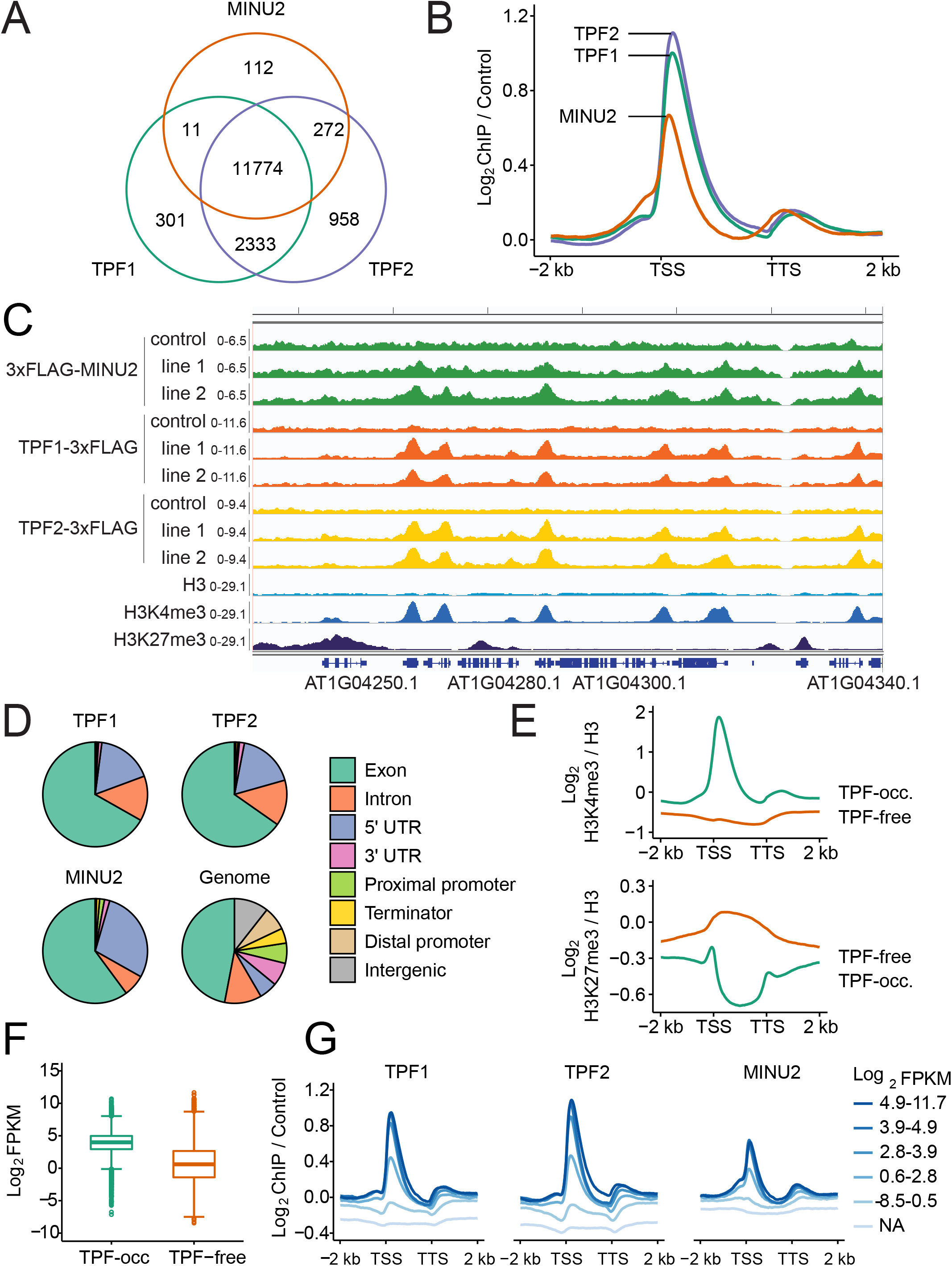
TPF and MINU2 proteins locate over the 5’regions of thousands of genes. A) Overlap between TPF1, TPF2, and MINU2 peaks. The peaks of each subunit are the union set of peaks detected (MACS2 q < 0.05) in ChIP-seqs from two independent transgenic lines. B) Average occupancy of TPF1, TPF2, and MINU2 over the intersection of TPF1 and TPF2 target genes (n = 15859), hereafter referred to as TPF-occupied genes. C) Browser screenshot showing the reproducibility of TPF and MINU2 peaks between lines. Notice these peaks co-localize with H3K4me3 and are absent in H3K27me3-enriched regions. Coverage values are Reads Per Million (RPM) mapped reads. D) Percentage of TPF1, TPF2, and MINU2 peak summits overlapping genomic features. Proximal promoters and terminators were defined as regions 500 bp upstream or downstream of the TSS or TTS, respectively. Distal promoters ranged from 500 bp to 2000 bp upstream of the TSS. Other non-genic regions were classified as intergenic. E-F) Average occupancy of H3K4me3 and H3K27me3 (E), and gene expression (F) over genes occupied by TPF (n = 15859) and genes lacking TPF peaks (hereafter TPF-free genes, n = 21574). G) Average occupancy of TPF and MINU2 over genes ranked by their expression level. The lowest group consists of nonexpressed genes (0 FPKM), while expressed genes were grouped into five quintiles of ascending expression.

### The plant BAF-like complex controls the +1 nucleosome positioning of thousands of genes

In order to investigate the role of the plant BAF-like complex on the genome-wide nucleosome landscape and study the contribution of TPF subunits to the complex’s activity, we performed MNase-seq experiments comparing the *minu1/2,* and the strong *tpf1/2* mutants (from now on *minu* and *tpf* mutants, respectively) with WT plants (Supplementary Data 6). This technique relies on the function of the Micrococcal Nuclease (MNase), which has endo-exonuclease activity and degrades faster the DNA that is not associated with nucleosomes, thus allowing the inference of the nucleosome positioning and the differences between nucleosomes in mutant and WT backgrounds. The MNase-seq replicates were highly reproducible (Supplementary Figure 8A,B). Moreover, the WT samples were very similar to a previously published MNAse-seq experiment using chromatin from the same tissue (Supplementary Figure 8B,C) (66). The changes observed in nucleosome position can be separated into changes in position, fuzziness, and occupancy (Figure 3A). Both the *tpf* and *minu* mutants presented changes in all three events but there was a clear preference for the control of positioning (Figure 3A, Supplementary Figure 9A,B). A visual inspection of the MNase-seq tracks revealed an overall shift of the +1 nucleosome towards the NDR, which was further confirmed with metaplot and heatmap representations over TPF target genes (Figure 3B,C,D, Supplementary Figure 9A, Supplementary Data 8). Importantly, this trend was not observed over non-target genes suggesting a direct implication of the plant BAF-like complex (Figure 3C,E). The distance between nucleosomes over target loci was not affected suggesting that this complex does not play a major role in nucleosome phasing. The shift observed was variable among genes and, on average, represented a 10-15 bp shift although some loci showed a consistent shift of more than 40 bp towards the NDR (Figure 3D,E). In summary, this data shows that the plant BAF-like complex is mainly involved in the control of the nucleosome positioning and prevents +1 nucleosome shift towards upstream regions.

**Figure 3.**
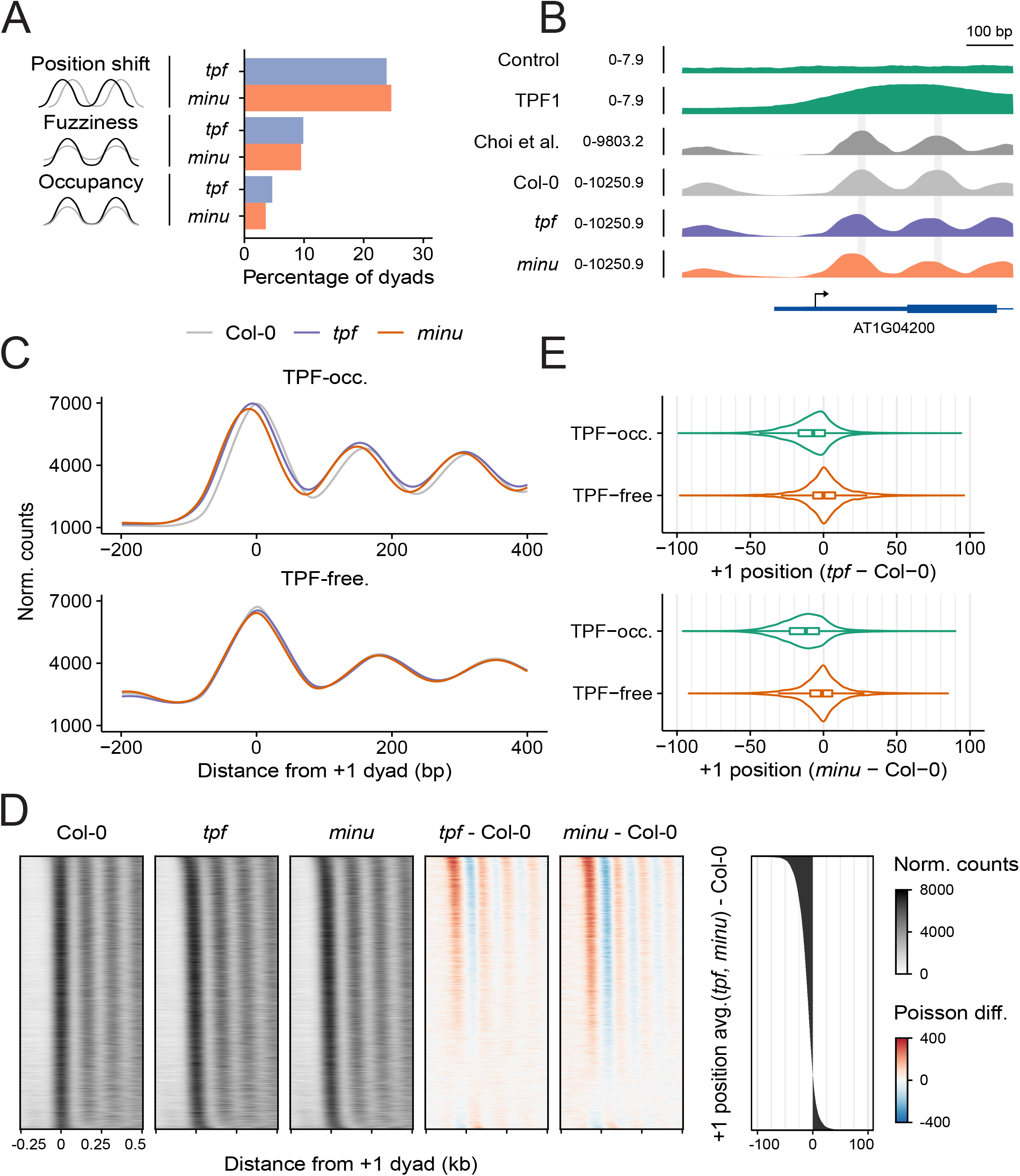
TPF and MINU control the position of the +1 nucleosome. A) Cartoon depicting the three different nucleosome features analyzed. Percentage of dyads showing significant position shift (|mutant – Col-0 position| > 5 bp), fuzziness (FDR < 0.05) and occupancy (FDR < 0.05) differences in *tpf* and *minu* mutants compared to Col-0. B) Representative TPF-occupied locus in which dyads are shifted towards the TSS in *tpf* and *minu* compared to Col-0. In green, ChIP-seq signal from Col-0 control and the mean Reads Per Million (RPM) of TPF1 ChIPs performed with two independent transgenic lines. In dark grey, nucleosome coverage from an independent study (66). Light grey, blue and orange represent normalized nucleosome coverages of Col-0, *tpf*, and *minu* mutants, respectively, from the mean of two biological replicates. In these loci, the +1 nucleosome is found 14 bp (*tpf*) and 23 bp (*minu*) upstream to the position of the Col-0 +1 nucleosome, scale bar 100 bp. C) Average nucleosome occupancy in Col-0, *tpf,* and *minu* mutants over TPF-occupied (n =11768) and TPF-free (n = 8891) genes that display well-positioned +1 nucleosomes. D) Heatmaps representing nucleosome positions in Col-0, *tpf,* and *minu* mutants ranked by the average +1 dyad shift observed in mutants. The first three heatmaps show normalized nucleosome occupancies of Col-0, *tpf,* and *minu* (average from two replicates), while the last two represent the poisson difference between Col-0 and mutant occupancies. The far-right plot shows the average +1 position shift between *tpf* and *minu* mutants ordered from most upstream to most downstream. E) Distribution of the +1 dyad shift values in *tpf* and *minu* mutants compared to Col-0 over TPF-occupied and TPF-free genes with well-positioned nucleosomes.

### The plant BAF-like complex affects the 5’ transcript length determination of thousands of genes with no significant effect on differential gene expression

Previous studies in yeast have shown that +1 nucleosome shift results in overall gene repression due to interference with the accessibility of the transcriptional machinery to the promoter (15,16,20). Thus, we performed RNA-seq experiments comparing the *tpf* and *minu* mutants with WT plants (Supplementary Figure 10A, Supplementary Data 6). Consistent with the strong developmental phenotype of these mutants, we found thousands of genes differentially expressed (DEGs), of which 2436 and 1225 were upregulated and 4081 and 2663 were downregulated in the *tpf* and *minu* mutants, respectively (Figure 4A, Supplementary Data 9). As expected, the DEGs from the two mutants showed a high degree of overlap although *tpf* mutant showed a larger number of up- and down-DEGs (Figure 4B, Supplementary Figure 10B). This is consistent with the stronger developmental phenotype of the *tpf* mutant compared to *minu*, which is a weak combination of *minu1* and *minu2* alleles (73) (Supplementary Figure 2D and 4). Importantly, the combination of strong *minu1 minu2* alleles causes embryo lethality, which hampers the ability to work with such material (73). In line with these results, both mutants showed similar enriched gene ontology categories, which were especially related to reproductive processes and response to biotic and abiotic stresses (Supplementary Figure 10C). Interestingly, we found that multiple subunits that are specific to the plant BAF-like complex were upregulated in the *tpf* and *minu* mutants while subunits specific to the plant ncBAF-like complex were not (Supplementary Figure 10D). This suggests that there is a negative feedback mechanism to upregulate the expression of the BAF-like-specific subunits in situations where the complex is not functional or is highly demanded.

**Figure 4.**
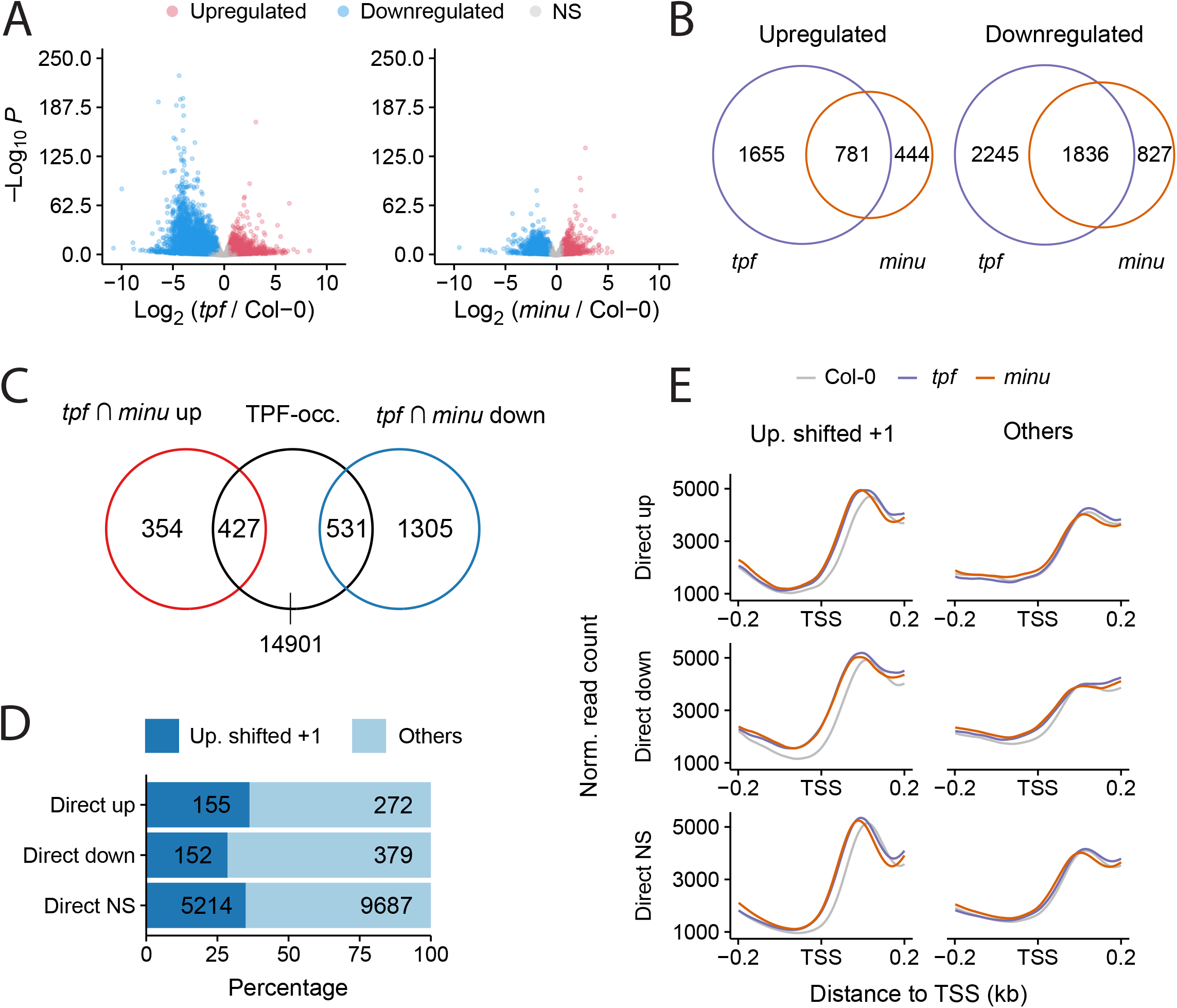
Changes in nucleosome positioning have a small impact on the differential expression of BAF-like targets. A) Volcano plots showing log2 differential expression of *tpf* and *minu* mutants compared to Col-0. Genes with P < 0.05 and fold change > 1.5 were considered differentially expressed. NS, not significant. B) Overlap between upregulated and downregulated DEGs detected in *tpf* and *minu* mutants. C) Overlap between TPF-occupied genes and shared DEGs of *tpf* and *minu* mutants. DEGs overlapping TPF-occupied genes were classified as either “direct up” or “direct down” depending on their misregulation direction. The rest were considered direct not significant (direct NS). D) Percentage of direct up, direct down or direct NS genes showing an upstream shift at the +1 dyad of at least 5 bp in both *tpf* and *minu* mutants compared to Col-0. E) Nucleosome occupancy profiles of direct up, direct down, and direct NS genes in Col-0, *tpf,* and *minu* mutants, centered on the TSS. Plots represent occupancy in loci that show a +1 nucleosome shift of at least 5 bp in both *tpf* and *minu* mutants compared to Col-0, or loci that do not show a shift (others). Occupancy values are normalized read counts averaged from two replicates.

In order to identify genes that are potentially regulated by the plant BAF-like complex, we selected TPF-occupied genes (which result from the intersection between TPF1 and TPF2 target genes) and looked for their overlap with DEGs. To work with highly confident BAF-like-regulated genes, we used the list of DEGs commonly misregulated in *tpf* and *minu* mutants (Figure 4B). We identified 427 direct targets that were upregulated and 531 targets that were downregulated, which together represented 6% of all direct targets (Figure 4C, Supplementary Data 10). This result indicates that most complex targets are not misregulated under the conditions where samples were taken. Perhaps sampling under more dynamic conditions would reveal a stronger effect. The number of DEGs that showed a +1 nucleosome shift was around one-third of the total number of direct DEGs (155/427 for up-DEGs and 152/531 for down-DEGs) but this trend was also shared by targets that were not differentially expressed, suggesting a small correlation between the +1 nucleosome shift and differential expression (Figure 4D). We compared the nucleosome profile over direct-up-DEGs, direct-down-DEGs, and direct-non-DEGs (Figure 4E). Interestingly, we found that the nucleosomal signal over the NDR region of the direct-down-DEGs was higher compared to the other two groups. This suggests that nucleosome invasion of the NDR over this set of genes could be responsible for the observed transcriptional downregulation in the *tpf* and *minu* mutants probably by interfering with the recruitment of the transcriptional machinery (15) (Figure 4E).

Remarkably, looking at the RNA-seq tracks we found a consistent increase in the accumulation of reads over the 5’ of many genes, suggesting an upstream change in TSS usage in the *tpf* and *minu* mutants (Figure 5A,B, and C). Importantly, the accumulation of these additional reads was only observed in genes targeted by the complex, suggesting that it may be participating in the TSS change (Figure 5B,C, Supplementary Figure 11A). This change in 5’ transcript length did not have an overall impact on gene expression since a similar 5’ differential enrichment was found in all TPF-occupied genes (Supplementary Figure 11B), of which only a fraction was differentially expressed (Figure 4C, Supplementary Figure 11B). Moreover, the accumulation of 5’ reads in *tpf* and *minu* mutants showed a remarkable correlation with their +1 nucleosome shift (Figure 5B,C, Supplementary Figure 11C), thus pointing to a functional connection between the 5’ transcript length determination and the +1 nucleosome repositioning. To validate the accumulation of the 5’ reads and confirm that they correspond to full-length transcripts, we studied 4 genes that showed this phenomenon and that were not differentially expressed (Supplementary Figure 11D). Results confirmed an increased accumulation of longer full-length transcripts in the *tpf* and *minu* mutants compared to Col-0 while overall expression levels were not affected (Figure 5D, Supplementary Data 11). These results indicate that the plant BAF-like complex plays an important role in determining the 5’ transcript length of its targets without affecting overall expression levels and also suggest that the +1 nucleosome repositioning could be functionally involved in such change of transcript length.

**Figure 5.**
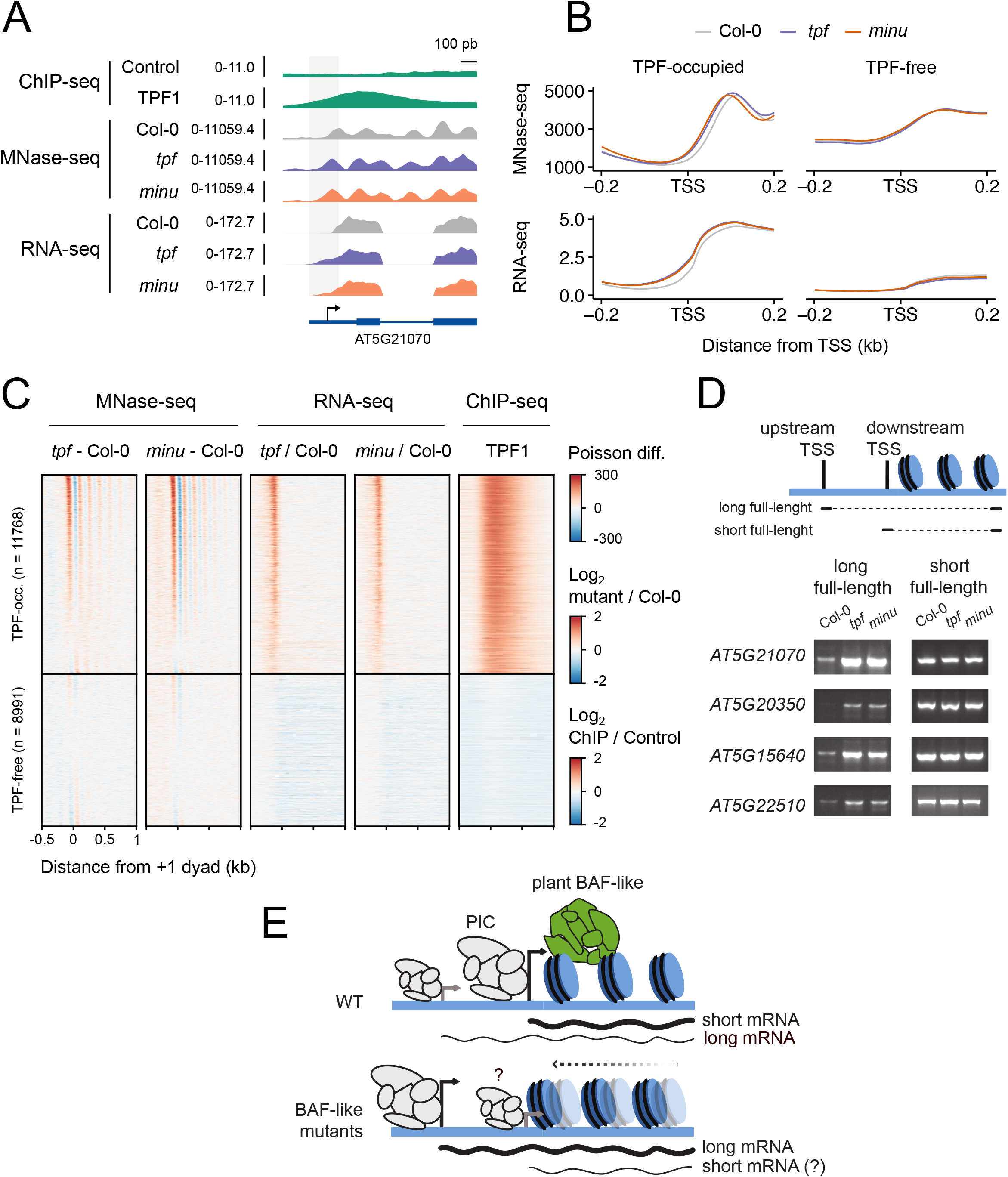
TPF and MINU affect the 5’ transcript length determination of thousands of genes. A) Example of a TPF-bound locus showing longer 5’ transcript length and +1 nucleosome shift towards the promoter in *tpf* and *minu* mutants compared to Col-0. ChIP-seq values are the mean Reads Per Million (RPM) of ChIPs performed with two independent transgenic lines. MNase-seq and RNA-seq values are normalized counts averaged from two and three biological replicates, respectively. B) Normalized nucleosome (upper part) and transcript coverage (lower part, shown in log2 scale) in Col-0, *tpf* and *minu* mutants over TPF-occupied and TPF-free genes. C) Heatmap showing MNase-seq poisson differences between genes with a properly positioned +1 nucleosome, RNA-seq log2 fold changes, and TPF1 ChIP-seq log2ratio values between transgenic lines and non-transgenic Col-0 controls. D) Expression of long and short full-length mRNAs of four selected genes that showed extended 5’ signal in the RNA-seq experiment in Col-0, *tpf* and *minu* mutants. The cartoon depicts the position of the primers used for each fragment. Since the accurate position of the TSS for the long isoform in the mutant backgrounds is not known, the Forward primers were chosen based on the reads from the RNA-seq experiment. The reverse primers were designed to bind to the region near the STOP codon. Primer information can be found in Supplementary Data 1. Amplicon sizes are (long/short; bp): AT5G21070 (937/831), AT5G20350 (2088/1939), AT5G15640 (1307/1100), At5G22510 (2328/1929). All bands were confirmed by Sanger sequencing. E) Proposed model of the role of the plant BAF-like complex on +1 nucleosome position and TSS usage. In WT plants, a dominant TSS drives the expression of the main mRNA isoform found in WT. In the *tpf* and *minu* mutants, a shift of the +1 nucleosome over the dominant TSS negatively affects its usage and promotes the use of an alternative upstream TSS resulting in the longer mRNA isoforms found in the *tpf* and *minu* mutants. Whether the shorter isoform is produced in the mutant backgrounds is not known at this point. The plant BAF-like complex is depicted in green while the Preinitiation complex (PIC) is depicted in grey. The size of the PIC cartoon represents its relative activity on the promoter.

## DISCUSSION

We recently identified an evolutionarily conserved plant SWI/SNF complex reminiscent of the mammalian BAF subclass (35). In the present study we report the function of the plant BAF-like complex, which regulates the position of the +1 nucleosome of thousands of genes and affects their 5’ transcript length. Moreover, we report experimental evidence that establishes the functional relevance of TPF proteins for the complex activity based on the similarities between the *tpf* and *minu* mutants.

### TPF proteins play an important role in the function of the plant BAF-like complex

Our results show that double *tpf* mutants present developmental phenotypes similar to the ones reported for *minu* mutants (73). Moreover, *tpf* and *minu* mutants show similar molecular phenotypes, including changes in gene expression, +1 nucleosome shift, and change in 5’ transcript length. This evidence strongly suggests that TPF proteins play a prominent role in the overall complex function. The characterization of the genome-wide TPF location shows a very high overlap with MINU2, reinforcing the idea that they are core components of the same complex. The fact that strong *minu* alleles cause more severe developmental defects than strong *tpf* alleles can be interpreted in two ways: either the BAF-like complex retains some activity in the absence of TPF, or the strongest *tpf* alleles used in this study are not null. The TPF orthologs in mammals participate in the recruitment of the complex to the chromatin through their tandem PHD domains that can recognize acetylated histones (49,50). The strong similarities between *tpf* and *minu* mutants suggest that TPF could be playing a significant role in the recruitment of the complex to their targets through their PHD and Tudor histone-reader domains. Future functional studies will shed light on these aspects.

### The BAF-like complex partially overlaps with ncBAF-like components

Our results show that the BAF-like complex is found over the 5’ regions of thousands of expressed genes. This location is shared by other SWI/SNF chromatin remodeling proteins in animals, and fungi (74,75). Studies of the genomic location of the BRM and SYD ATPases revealed their location over TSS regions of many genes (38,39,76), pointing to their colocalization with the BAF-like complex. An open question is whether there is functional crosstalk between the plant BAF-like and ncBAF-like complexes over shared targets. Importantly, it remains to be shown whether the SYD ATPase is incorporated into a ncBAF-like complex or into an uncharacterized plant SWI/SNF subclass. However, previously reported functional redundancy between BRM and SYD (40) supports its presence in ncBAF-like. BRM and SYD ATPases were also found over more distal promoter regions and downstream genic regions (39,42,76). Indeed, BRM participates over the 3’ regions of genes to regulate antisense transcription (77). Our results did not show significant enrichment of the BAF-like complex over distal promoter and terminator regions indicating that the plant BAF-like and ncBAF-like complexes can bind to distinct targets to perform non-redundant functions. We found a strong correlation in the position of the BAF-like complex with permissive marks like H3K4me3 and H3panAc marks. The fact that TPF contains 3xPHD and Tudor domains, previously shown to interact with methylated and acetylated histones (49,50,78), makes these two histone marks likely direct targets of TPF proteins to promote recruitment or stability of the complex on target sites. On the contrary, the complex was not found over facultative or constitutive heterochromatic regions, similar to what was observed for BRM and SYD targets, indicating that plant SWI/SNF complexes do not normally localize to these regions (38,39,76). Interestingly, BRM and SYD have been shown to play antagonistic and synergistic interactions with the Polycomb complex, responsible for the deposition of the facultative heterochromatic mark H3K27me3 (79–81). Future studies will reveal if the BAF-like complex plays similar roles. The BAF-like complex contains the plant-specific SHH2 subunit able to interact with the constitutive heterochromatic mark H3K9me1 in *in vitro* assays (82,83). Where and when the H3K9me-reader ability of SHH2 becomes required in the BAF-like complex remains to be explored.

### The plant BAF-like complex controls the positioning of the +1 nucleosome at thousands of loci

Depletion of the plant BAF-like complex showed a genome-wide effect on the positioning of the +1 nucleosome, with smaller effects on the fuzziness and occupancy of nucleosomes. A similar +1 nucleosome shift has been reported in yeast SWI/SNF mutants indicating a functional conservation of their activity (15,20). However, the distance between nucleosomes was not altered indicating that the plant BAF-like complex is not required to control nucleosome phasing, which has been attributed to the ISWI remodeler in plants (84). This is consistent with results in yeast where RSC depletion triggered +1 nucleosome shift with no effect on internucleosomal spacing (85,86). A recent study on the genome-wide impact of BRM depletion on nucleosomes reported defects mostly related to the control of the occupancy of nucleosomes flanking the BRM targets, especially those modified with the H2A.Z variant (38). This indicates that plant BAF-like and ncBAF-like complexes can play unique roles in setting up the nucleosomal landscape. Importantly, while there is information about BRM (38) and MINU (this study), no study so far has reported the genome-wide impact on nucleosomes of the remaining ATPase in plants –SYD-, which would provide a complete picture of the effect of the different plant SWI/SNF ATPases on the nucleosome landscape.

A genome-wide analysis of the effect of plant BAF-like depletion on the transcriptome revealed that only a small number of direct targets were misregulated. This is despite the large number of direct targets that show a +1 nucleosome shift, suggesting that this change in nucleosome positioning does not play a prominent role in the regulation of transcript levels. There are examples in fungi and animals where an upstream nucleosome shift towards the NDR occludes the dominant TSS and triggers a global downregulation of expression (16,19,20). While we did not see such global effect, we did observe a correlation between downregulated direct targets and a stronger nucleosome invasion of the NDR. Perhaps more dynamic situations such as during the response to environmental or developmental cues would reveal a larger number of genes whose expression is affected by the upstream repositioning of the +1 nucleosome.

### The plant BAF-like complex affects the 5’ transcript length determination of thousands of genes

Interestingly, we observed that thousands of plant BAF-like complex targets showed an extended 5’ region in the *tpf* and *minu* mutants pointing to the participation of the complex in the selection of alternative TSSs. This change in 5’ transcript length did not affect the overall expression level of most target genes suggesting that the promoter of these genes can still drive the same amount of expression regardless of the TSS used. Recent studies have reported that approximately 75% of genes are estimated to use multiple TSSs in Arabidopsis (87,88). Thus, the new TSSs used in the plant BAF-like mutants most likely correspond to previously found TSSs that are not preferentially used in unchallenged conditions but might be chosen in certain developmental or environmental situations. Importantly, changes in TSS selection have been reported in different tissues or upon environmental changes (87,89–91). The extended 5’ transcript regions observed in the *tpf* and *minu* mutants could influence mRNA function through, for example, the incorporation of upstream ORFs and localisation signals that can impact translation efficiency and final localisation of proteins, respectively (87,89). Perhaps these events could be responsible for some of the pleiotropic defects observed in the *tpf* and *minu* mutants.

We found a remarkable correlation between the changes in +1 nucleosome positioning and 5’ transcripts length observed in the *tpf* and *minu* mutants. Previous reports in yeast have proposed a role for the +1 nucleosome in determining the usage of the TSS (16,92,93). In yeast SWI/SNF mutants, an upstream nucleosome shift towards the NDR can position the TSS closer to the midpoint of the nucleosome, which is a more repressive location (74). Moreover, +1 nucleosome shifts can directly impact the ability of TATA-binding protein (TBP) to bind to the promoter hampering the use of the dominant TSS (15). Thus, in promoters with multiple TSSs, those not occluded by the shifted nucleosome would remain active and allow gene expression (16). Our results suggest that this could be happening in the *tpf* and *minu* mutants, in whom a +1 nucleosome shift towards the NDR could be reducing the use of the dominant TSS responsible for the main mRNA isoform found in WT plants. This would in turn promote the selection of upstream TSSs, resulting in the longer mRNA isoforms found in the *tpf* and *minu* mutants (Figure 5E). Interestingly, we observed a positive correlation between the upstream displacement of the +1 nucleosome shift and the amount of reads differentially accumulated in the 5’ transcript ends, suggesting that the new +1 nucleosome position could be dictating the position of the alternative TSS, as has been previously suggested in yeast studies (16,92,93).

In summary, this study reports the function of the plant BAF-like complex as an important regulator of the +1 nucleosome positioning and the 5’ transcript length determination and provides experimental evidence of the functional relevance of the TPF signature subunits. We envision a scenario where internal and external stimuli could modulate the activity of the plant BAF-like complex to influence the 5’ length of transcripts. This would result in altered mRNA functionality that could contribute to the adaptive response of the plant to the triggering stimuli. Future work should focus on several open questions such as which factors influence the final position of the +1 nucleosomes in the plant BAF-like mutants or what is the mechanistic connection between the +1 nucleosome shift and the selection of specific upstream TSSs.

## DATA AVAILABILITY

Data supporting the findings of this work are available within the article and its supplementary materials. The genomics data have been deposited to the GEO repository with accession number GSE205112.

## ACKNOWLEDGEMENT

We thank Drs Miguel Blázquez and Maria A. Nohales for critical reading and discussion of the manuscript.

## AUTHOR CONTRIBUTION

B.D-M. performed all the *tpf* phenotyping experiments. B.D-M., and J.G-B. performed ChIP-seq, IP-MS, and MNase-seq experiments. J.P-A. performed all bioinformatics analyses. B.D-M., J.C-F., J.P. performed the Y2H experiments. D.A. supervised the work and edited the manuscript. Y.J-A., and J.W. performed the IP-MS analyses. B.D-M., J.P-A., and J.G-B. conceived this study and wrote the manuscript.

