## Supplementary Figures for "The TRIPLE PHD FINGERS proteins are required for SWI/SNF complex-mediated +1 nucleosome positioning and 5’ transcript length determination in Arabidopsis"

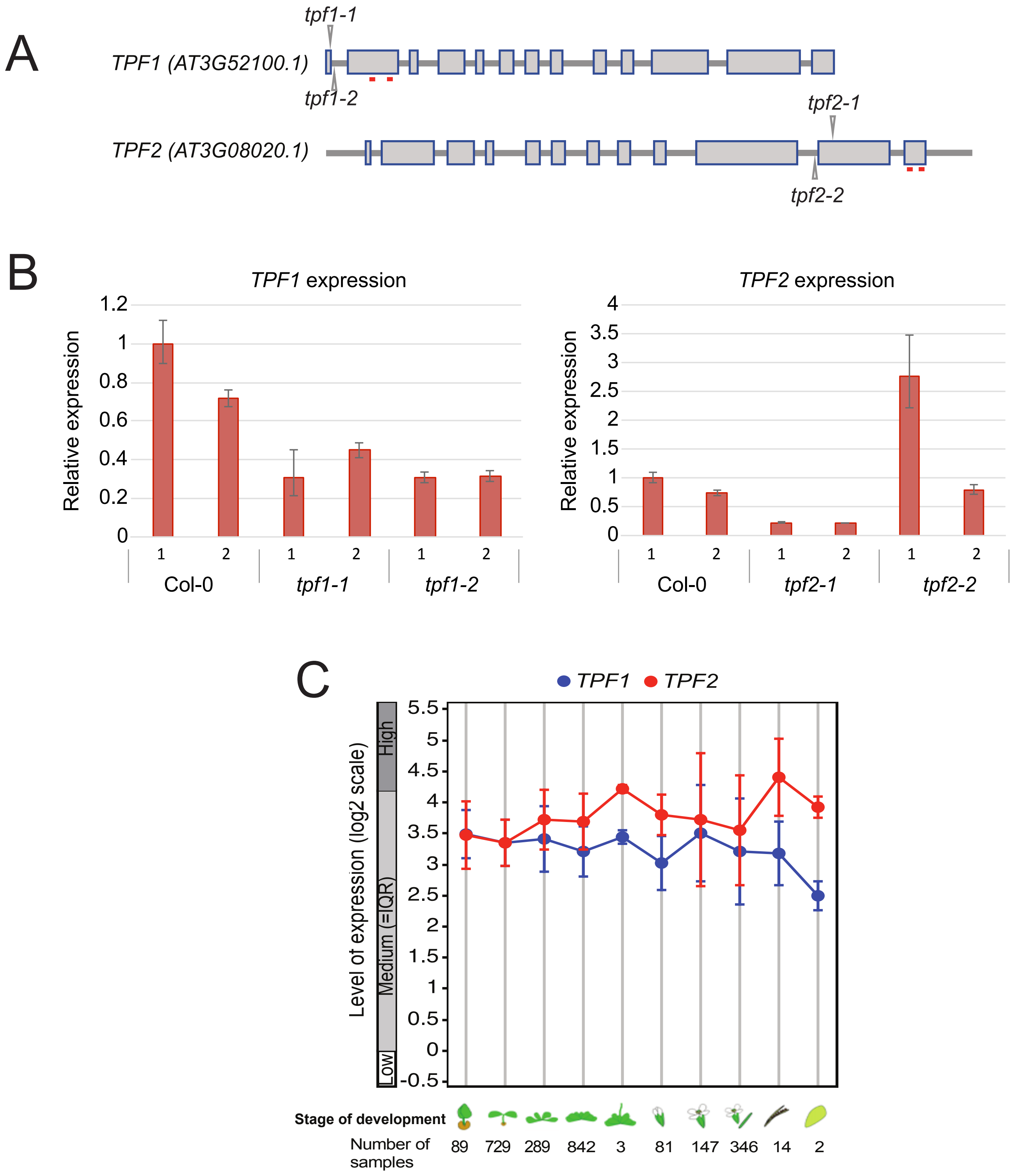


**Supplementary Figure 1. Molecular characterization of the *TPF* T-DNA insertion lines and expression levels.** A) Schematic representation of the T-DNA insertion sites (marked with triangles) in the genomic *TPF1* and *TPF2* loci. Location from the start codon: 30 bp (*tpf1-1*), 36 bp (*tpf1-2*), 3152 bp (*tpf2-1*), and 3050 bp (*tpf2-2*). Location of the primer pairs used to study the expression of the *TPF* genes in the T-DNA lines are highlighted as red squares. B) Relative expression of *TPF1* and *TPF2* normalized against *PP2A* in Col-0 plants and the T-DNA lines described in (A). Values of two biological replicates are shown. C) Expression levels of *TPF1* and *TPF2* at 10 developmental stages from AT_mRNASeq_ARABI_GL-3 dataset provided by Genevestigator.

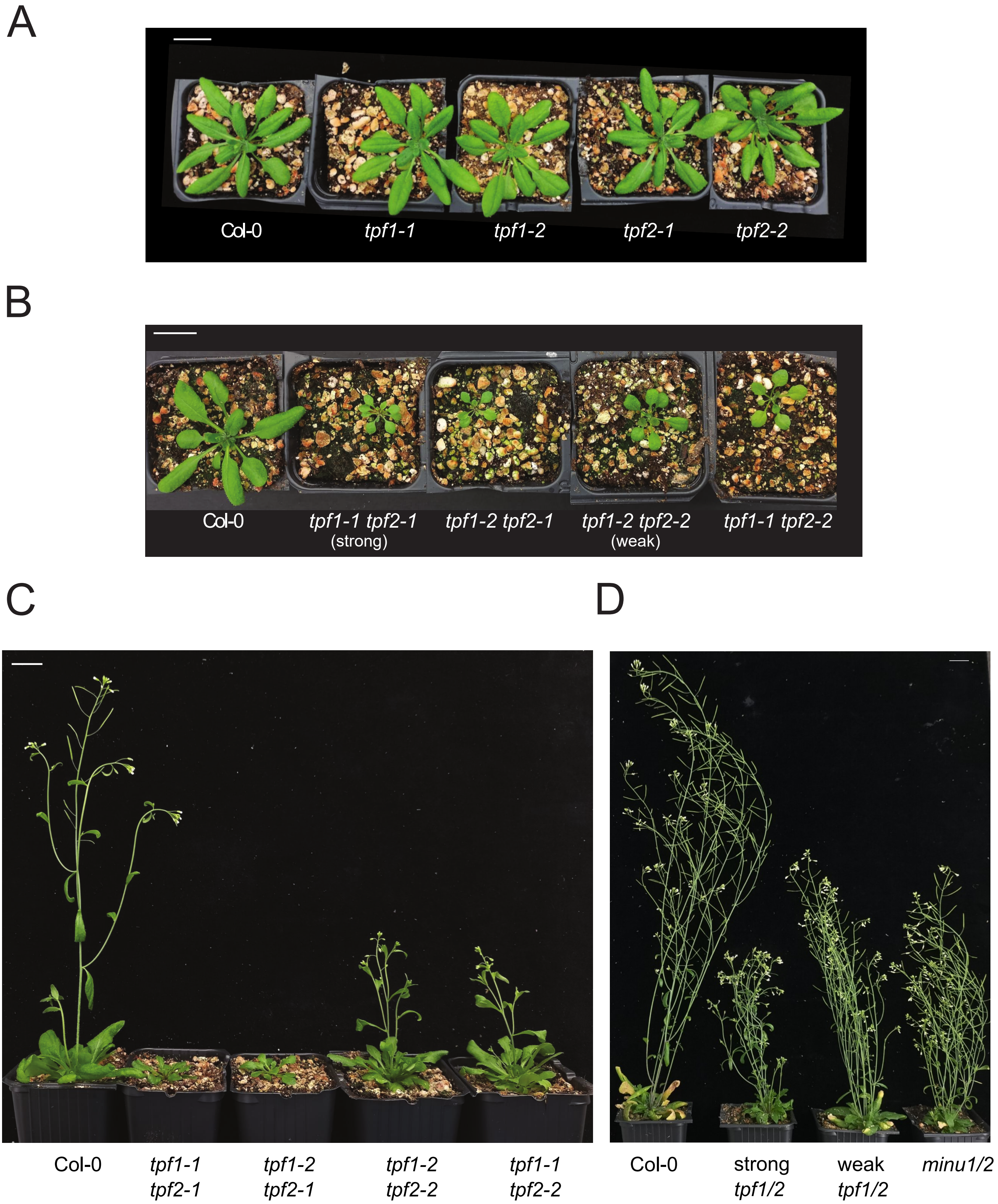


**Supplementary Figure 2. Phenotypical characterization of the single and double *tpf* mutants.** A) Four-week-old Col-0 and *tpf* single mutants grown on soil in long-day condition, scale bar: 2 cm. B-C) Three- and five-week-old *tpf1/2* allele combinations grown on soil in long-day condition, scale bar: 2 cm. D) The same image shown in Figure 1E including a representative plant of *minu1-2 minu2-1* (*minu1/2*) double mutant, scale bar: 2 cm.

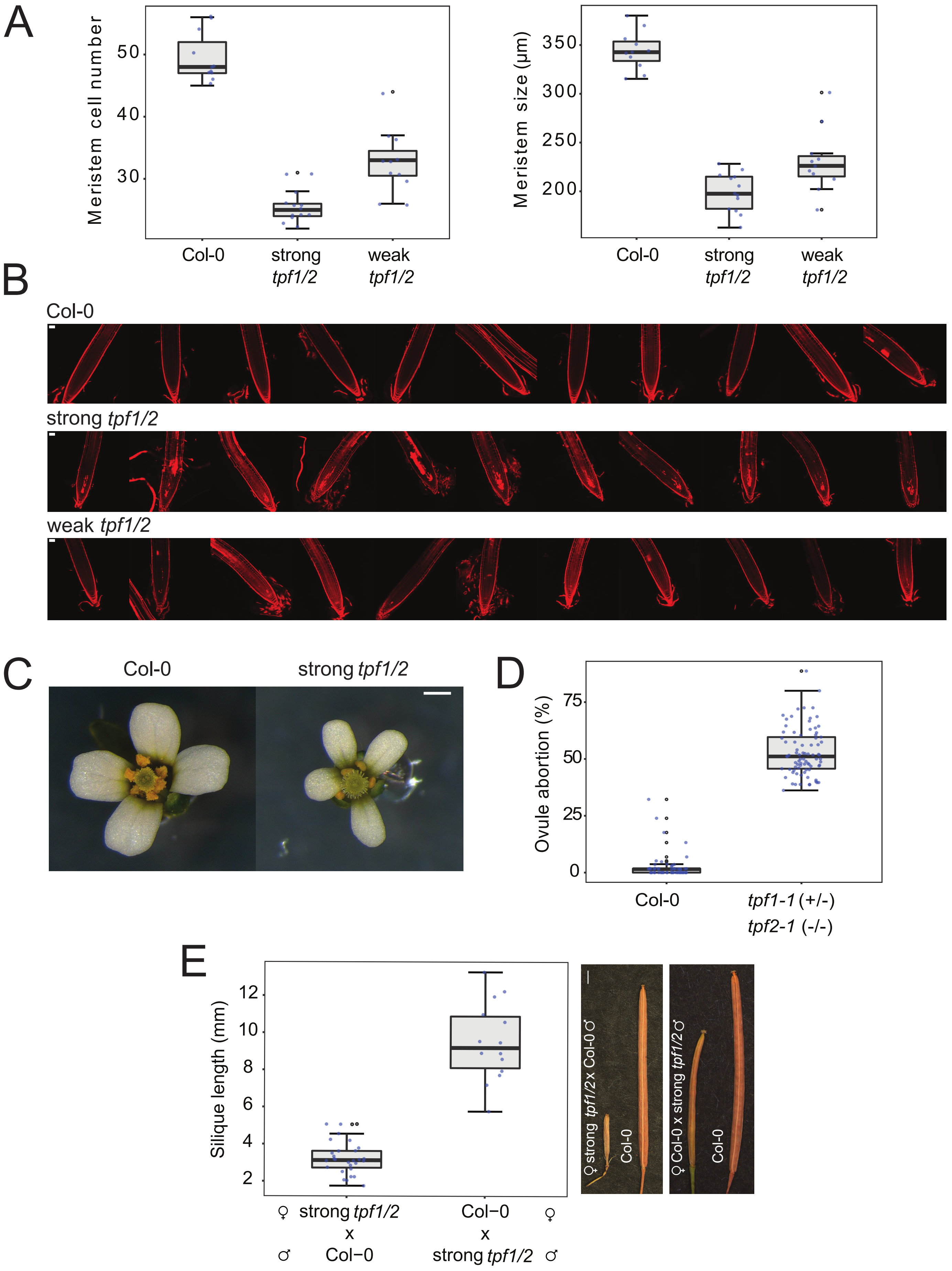


**Supplementary Figure 3. Phenotypical characterization of strong and weak *tpf* mutants.** A) Root meristem cell number and root meristem size from twelve-day-old in-vitro-grown seedlings of Col-0 (n=11), strong (n=13), and weak (n=11) *tpf1/2* mutants. The differences between genotypes are significant according to the one-way ANOVA-Bonferroni multiple comparisons test, P<0.05. B) Confocal microscopy images of eleven roots of twelve-day-old in-vitro-grown seedlings of Col-0, strong, and weak *tpf1/2* mutants stained with propidium iodide, scale 50 µm. C) Representative images of Col-0 and strong *tpf1/2* mutant flowers at anthesis, scale bar: 0.5 mm. D) Percentage of ovule abortion in the siliques corresponding to the 12-20 first positions of the principal stem of Col-0 (n=57) and *tpf1-1 (+/-) tpf2-1 (-/-)* (n=81) plants. E) Length of siliques resulting from reciprocal crosses between the strong *tpf1/2* mutant and Col-0 (left) (n=28, and n=14, respectively), and representative images of siliques from the reciprocal crosses and Col-0 plants (right), scale bar: 1 mm. Differences in D) and E) are statistically significant according to unpaired two-tailed t-tests, P-value < 0.05.

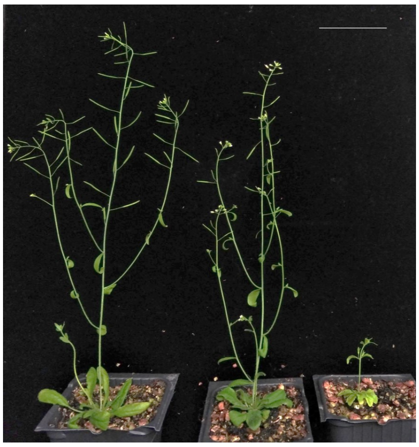

Col-0 TPF1-3xFLAG strong *tpf1/2* strong *tpf1/2*

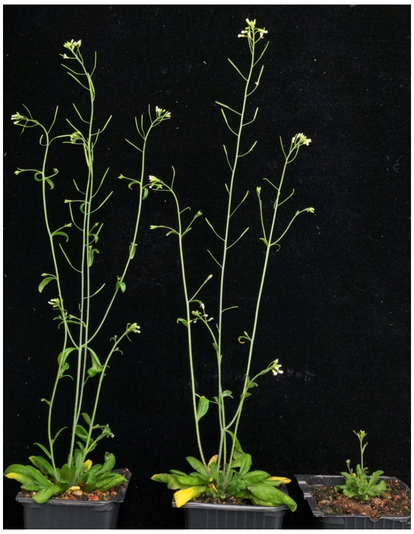

Col-0 TPF2-3xFLAG strong *tpf1/2* strong *tpf1/2*

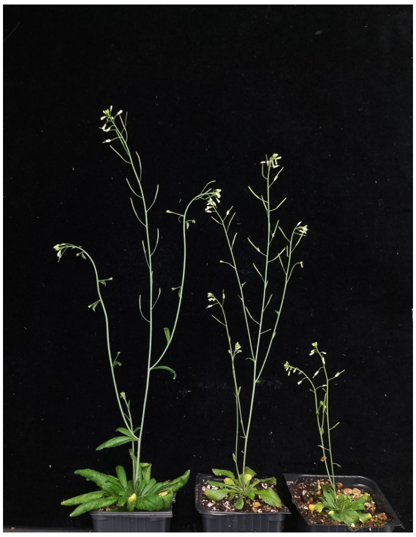

Col-0 3xFLAG-MINU2 *minu1/2* *minu1/2*

**Supplementary Figure 4. 3xFLAG-tagged *TPF* and *MINU2* transgenes can rescue the strong *tpf* and *minu* mutant phenotypes, respectively.** Plants expressing TPF1-3xFLAG or TPF2-3xFLAG in the strong *tpf1/2* mutant background, as well as expressing 3xFLAG-MINU2 in the *minu1/2* double mutant are shown. Corresponding untransformed controls (Col-0, strong *tpf1/2*, and *minu1/2* plants) are shown for comparison. Scale bar, 5 cm.

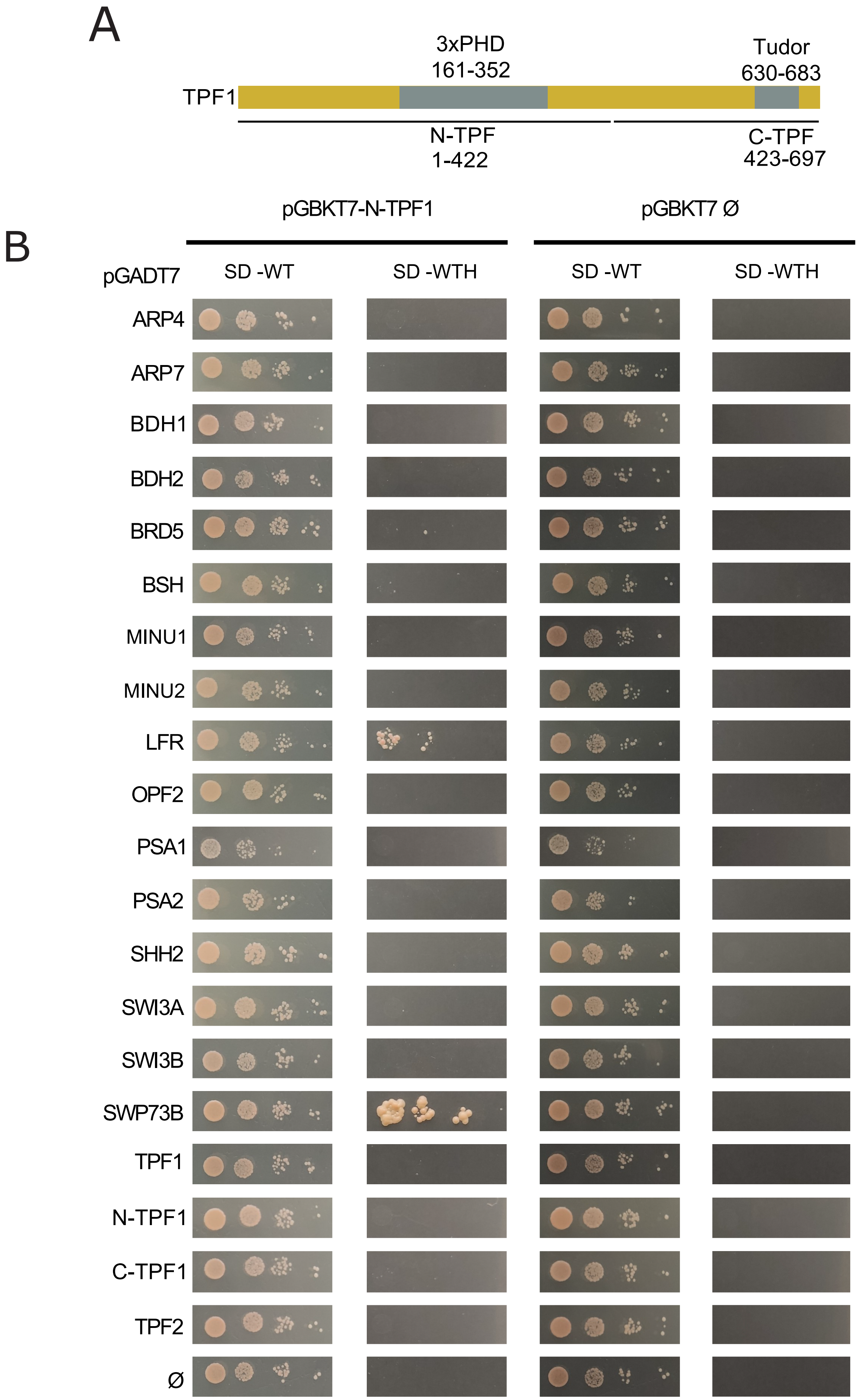


**Supplementary Figure 5. TPF1 interacts with SWP73B and LFR by Y2H.** A) Schematic representation of the TPF1 full-length protein depicting the amino acids corresponding to the 3xPHD and Tudor domains, as well as the N-TPF1 and C-TPF1 fragments. B) Yeast two-hybrid assay showing the interaction of N-TPF1 with SWP73B and LFR. L, Leucine; W, Tryptophan; H, Histidine and ∅, empty plasmid.

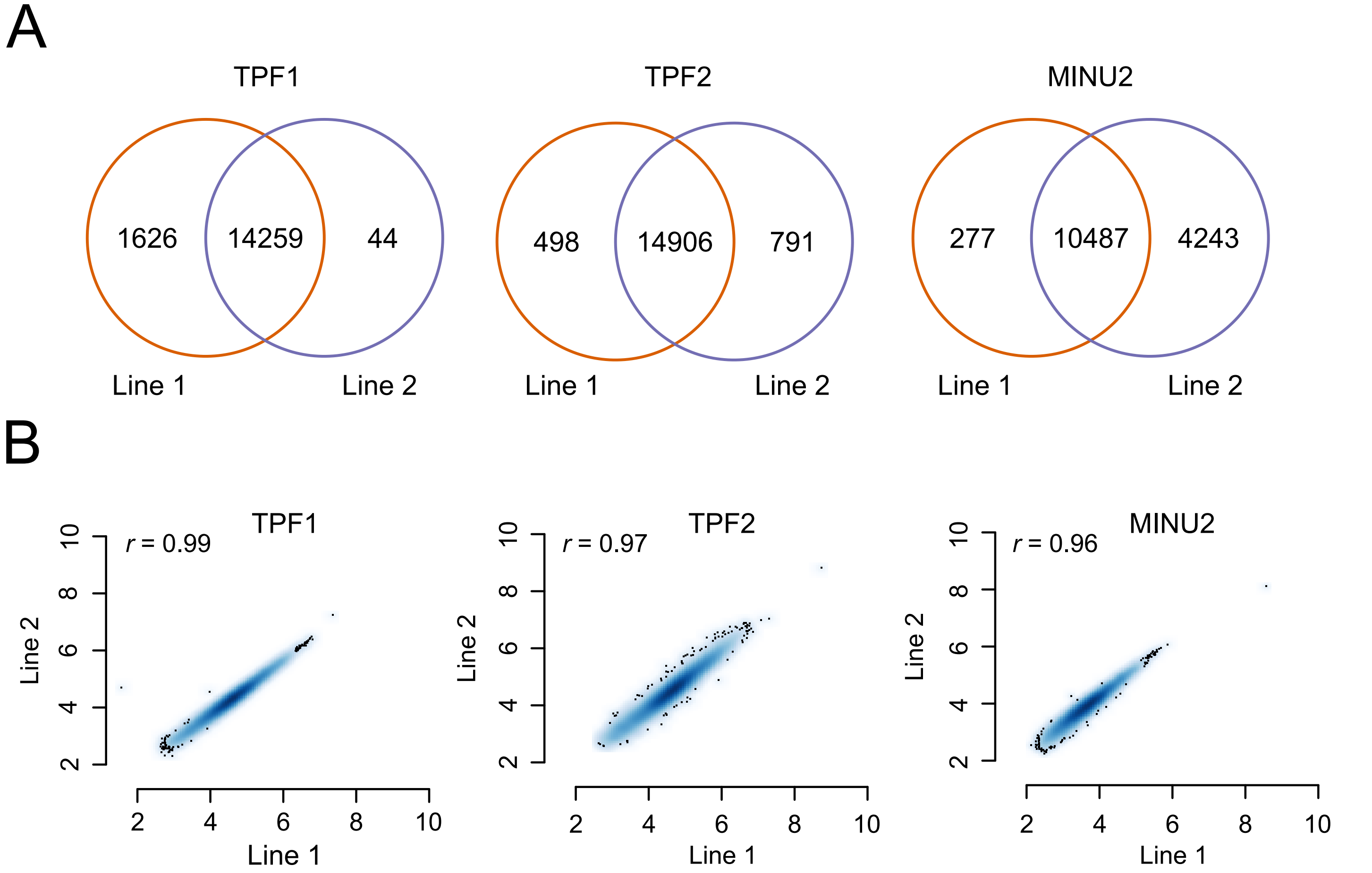


**Supplementary Figure 6. TPF and MINU ChIP-seqs are highly reproducible.** A) Overlap between TPF1, TPF2 and MINU2 peaks that were detected in ChIP-seq experiments involving two independent transgenic lines compared to Col-0 control (MACS2 $q < 0.05$). B) Density scatter plots showing the correlation (pearson coefficient) between read counts over peaks from TPF1, TPF2 and MINU2 lines. Reads were counted in the merged peak set of each experiment and normalized to Reads Per Million. Plot axes are in log2 scale.

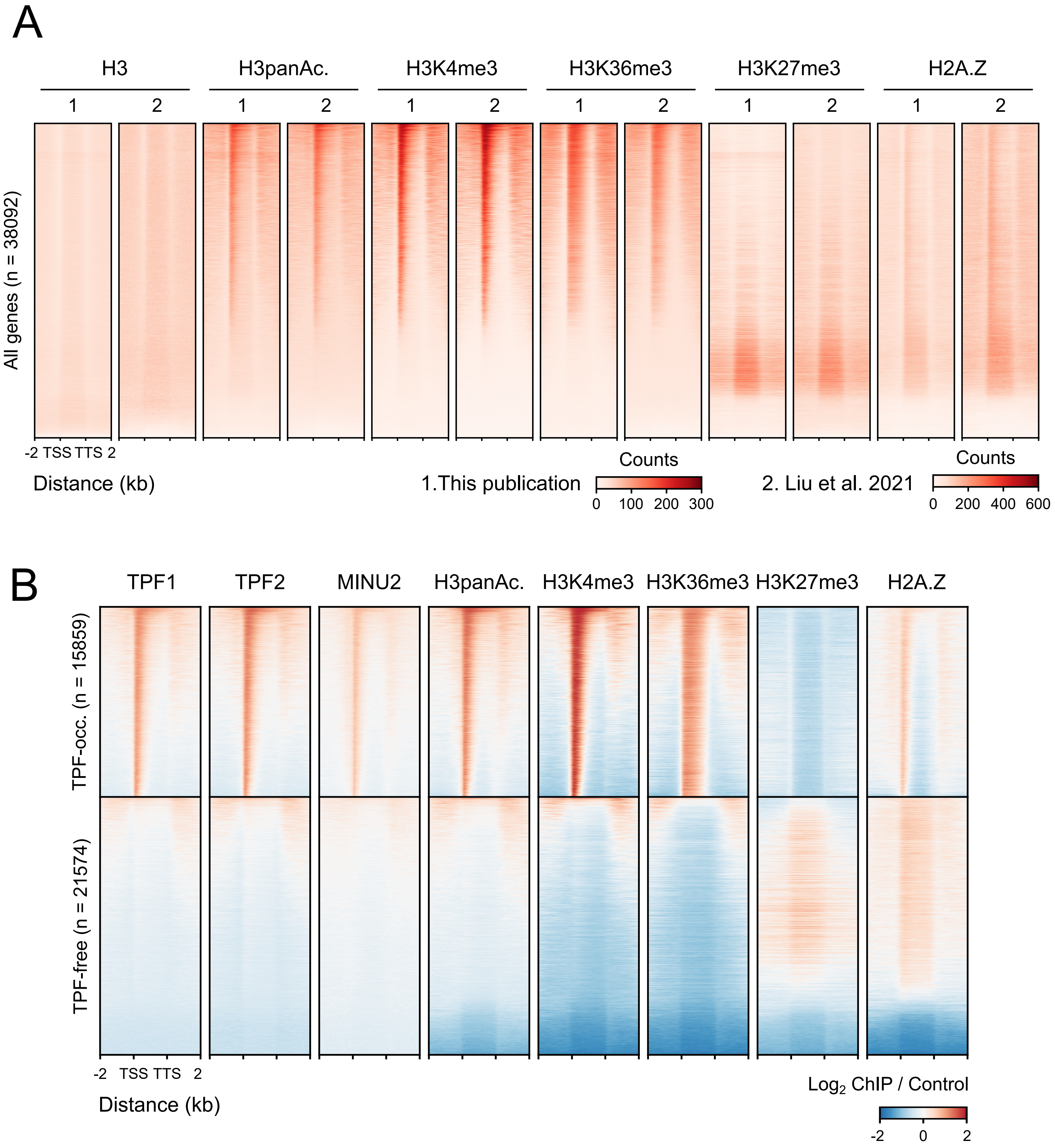


**Supplementary Figure 7. TPF and MINU2 co-localize with permissive epigenetic marks genome-wide.** A) Heatmaps comparing the accumulation of H3, H3panAc, H3K4me3, H3K36me3, H3K27me3, and H2A.Z derived from ChIP-seq data of this publication and a previous publication (63) over all the Araport11 genes. B) Heatmap showing the accumulation of TPF1, TPF2, MINU2, H3panAc, H3K4me3, H3K36me3, H3K27me3, and H2A.Z over TPF-occupied and TPF-free genes. TPF1, TPF2 and MINU2 occupancy values are the average of two independent transgenic lines and were log2-ratio normalized against untransformed Col-0 controls. H3panAc, H3K4me3, H3K36me3, H3K27me3, and H2A.Z occupancy values were log2 ratio-normalized against H3.

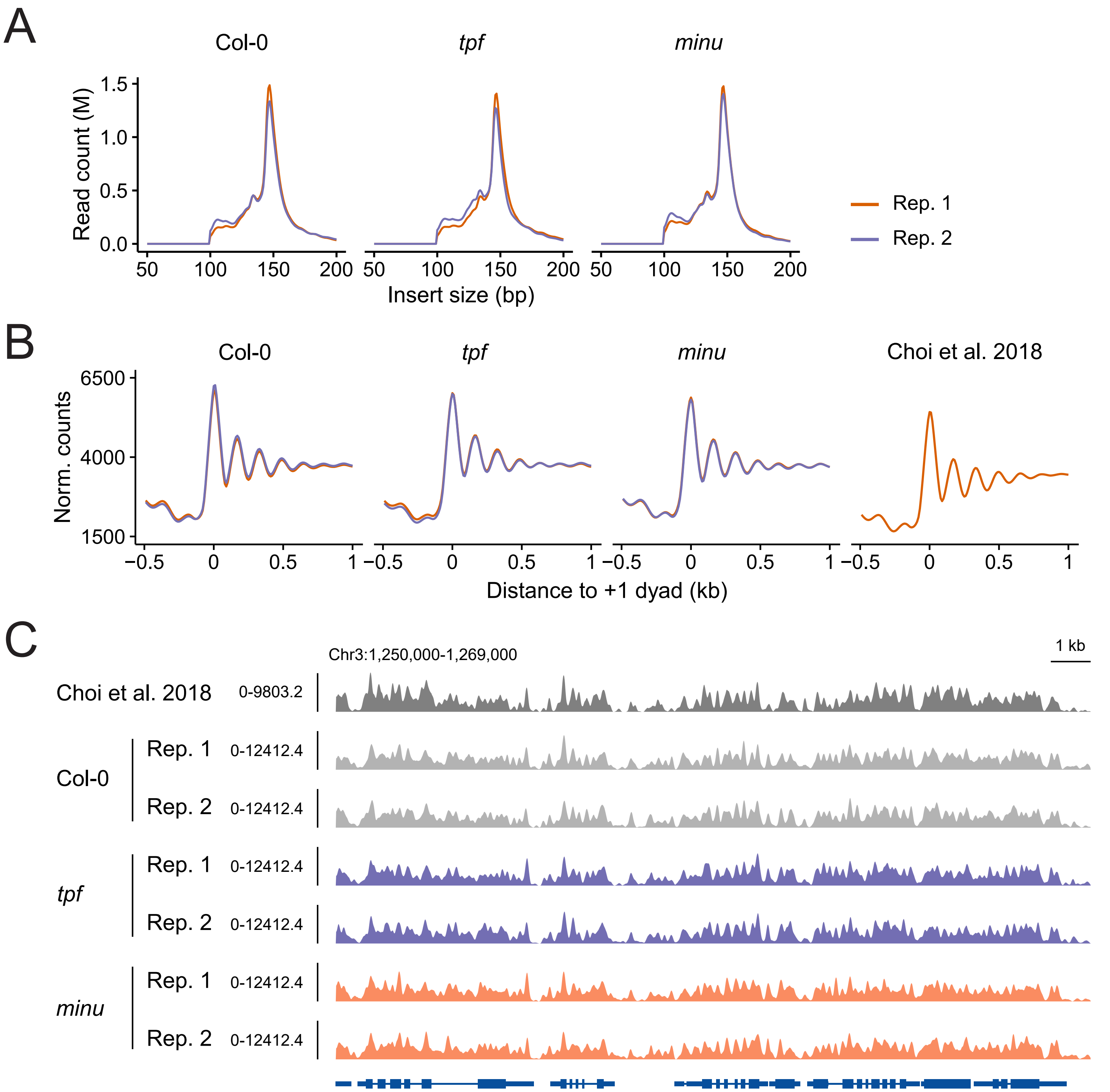


**Supplementary Figure 8. MNase-seq data is highly reproducible.** A) Insert size distributions of Col-0, *tpf* and *minu* MNase-seq replicates. The most frequent insert size in all samples is 147 bp as expected from MNase-seq data. B) Nucleosome occupancy profiles of Col-0, *tpf* and *minu* replicates together with Col-0 from a previous study (66). The plots were centered in the Col-0 +1 dyad positions obtained in this study. C) Genome browser screenshot displaying the nucleosome occupancy patterns of Col-0, *tpf* and *minu* replicates and Col-0 from (66). Values represent normalized read counts.

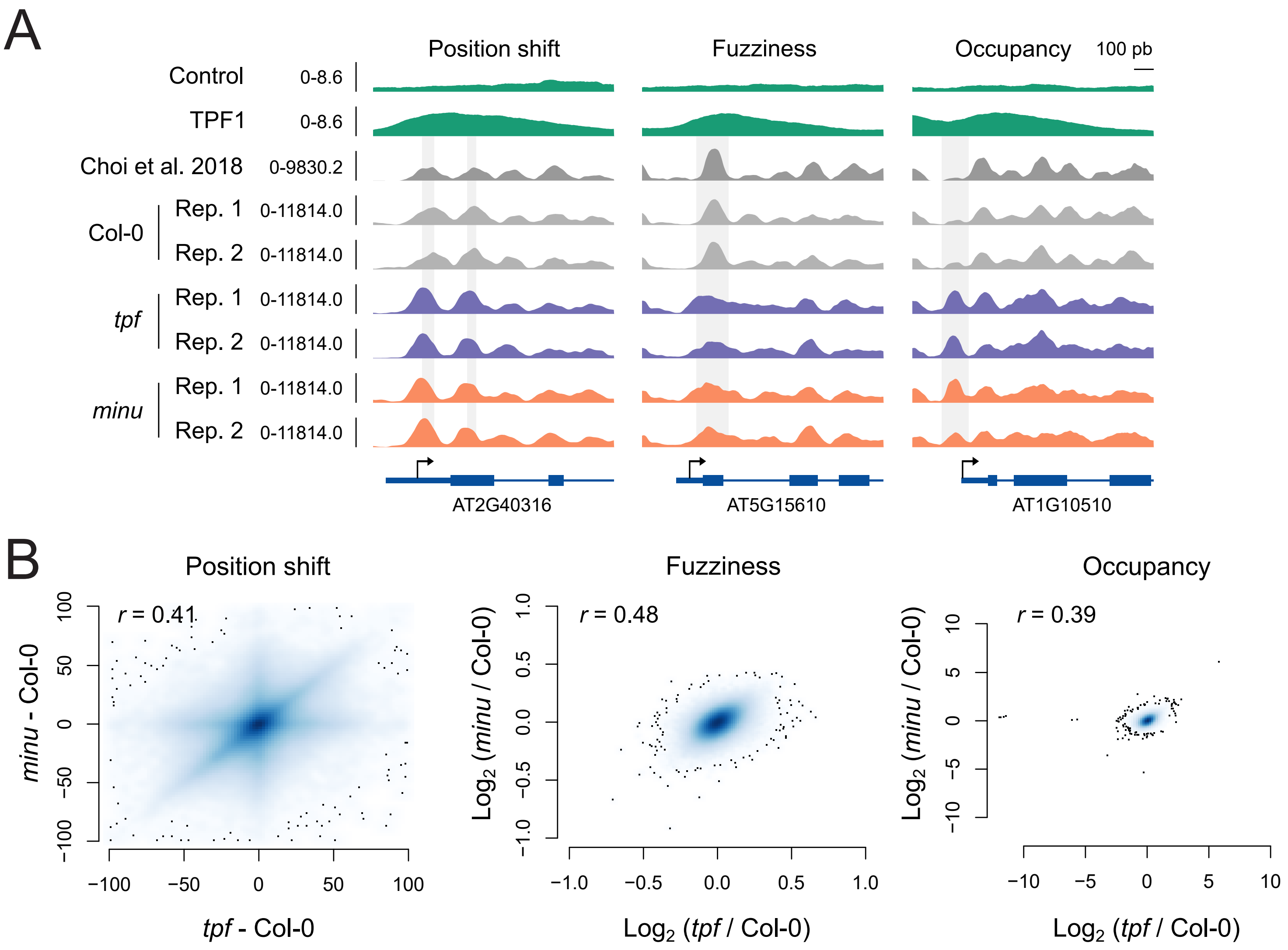


**Supplementary Figure 9. The impact of TPF and MINU on nucleosome positioning is reproducible across replicates and mutant backgrounds.** A) Examples of TPF-targeted loci that undergo different nucleosome changes in *tpf* and *minu* mutants. ChIP-seq values of TPF1 are normalized read counts averaged from two independent transgenic lines, while MNase-seq values are normalized read counts. Shaded areas depict the observed nucleosome changes. B) Density scatter plots showing the correlation (pearson coefficient) between position shift values, fuzziness fold changes, and occupancy fold changes obtained from comparing *tpf* and *minu* mutants to Col-0.

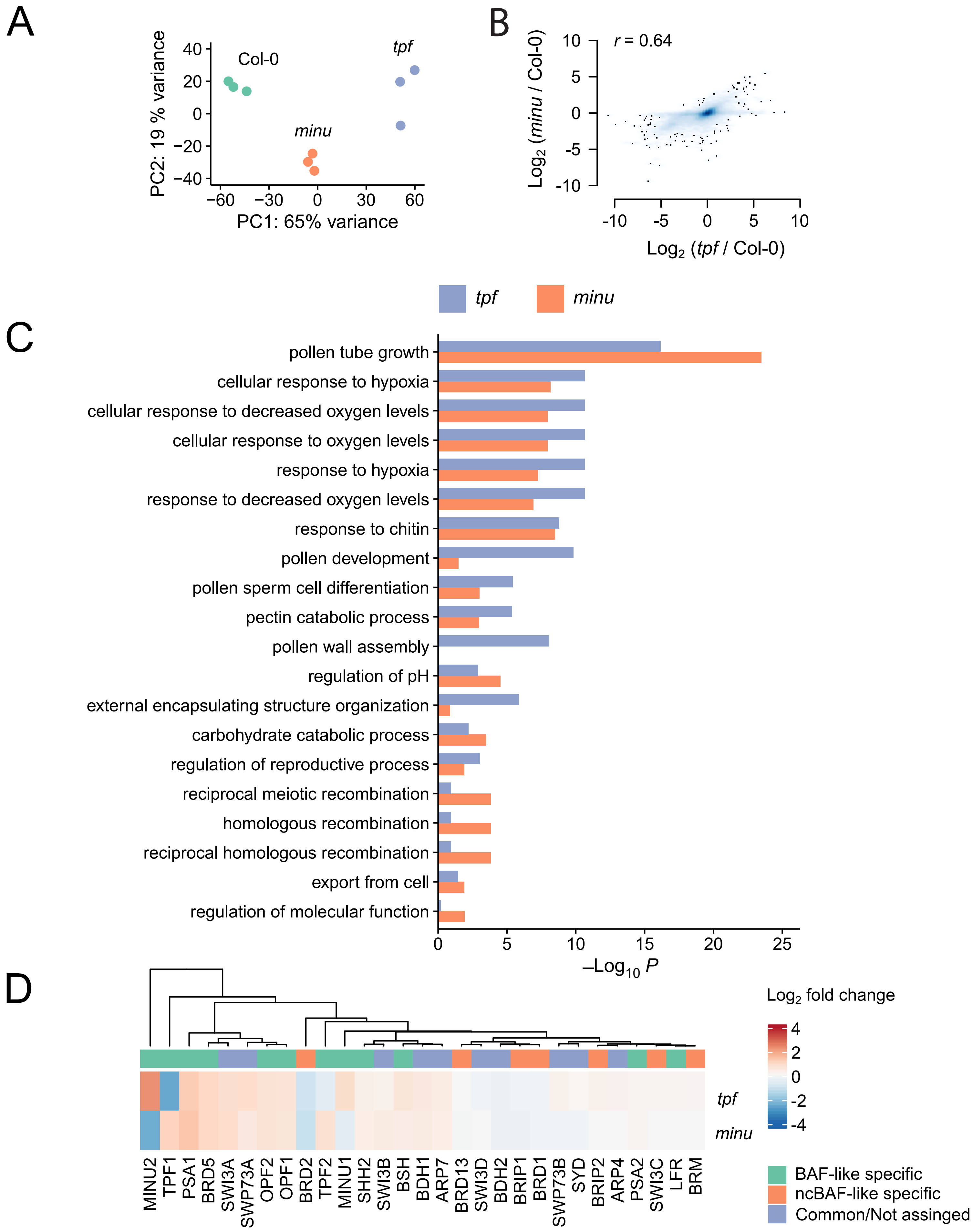


**Supplementary Figure 10. TPF and MINU regulate similar transcriptomes and gene ontology categories.** A) Principal Component Analysis performed over the transcriptome profiles of Col-0, *tpf*, and *minu* samples. The analysis was computed over DESeq2 r-log-transformed counts. B) Density scatter plot showing the correlation (pearson coefficient) between log2 fold changes of *tpf* and *minu* mutants from differential expression analyses against Col-0. C) Gene Ontology enrichment analysis over *tpf* and *minu* DEGs. The top twelve most significantly enriched terms from each analysis were selected. D) Heatmap showing log2 fold changes of known plant SWI/SNF subunits in the *tpf* and *minu* mutants compared to Col-0. Assignment of the different subunits to either BAF-like or ncBAF-like is based on experimental evidence from proteomics experiments from different studies (35). Subunits specific to the plant BAF-like complex are marked in green, those specific to the ncBAF-like complex are marked in orange, and those common to both complexes or where there is not enough evidence to assign to BAF/ncBAF are marked in blue.

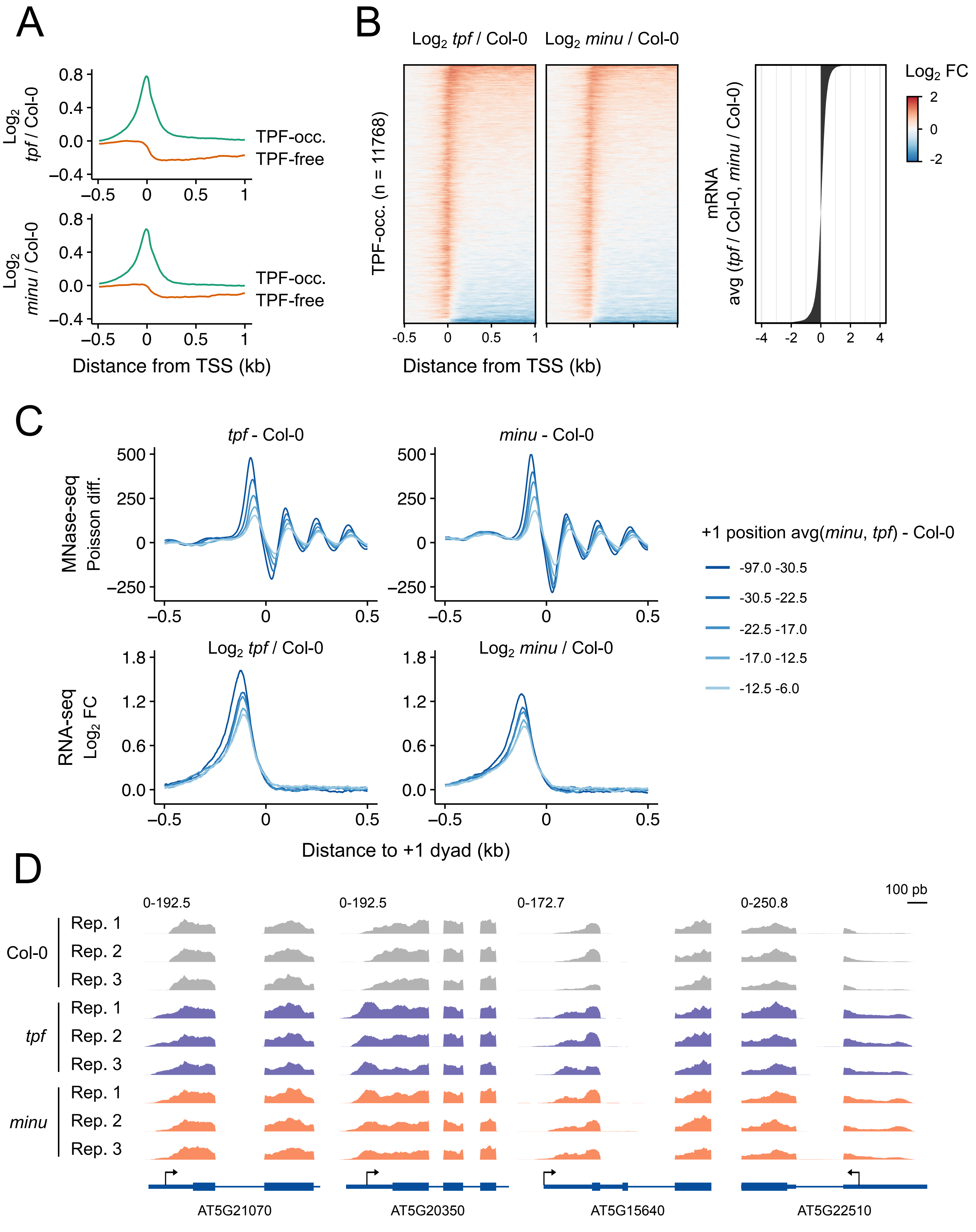


**Supplementary Figure 11. TPF and MINU regulate the 5´ transcript length determination**. A) Metaplots of RNA-seq log2 fold changes of *tpf* and *minu* mutants over TPF-occupied and TPF-free genes that exhibit a well-positioned +1 nucleosome. B) Heatmap showing RNA-seq log2 fold changes of *tpf* and *minu* over TPF-occupied genes ranked by the average log2 fold change of *tpf* and *minu* obtained by comparing their mRNA expression to Col-0. C) Metaplots showing the correlation between change in +1 nucleosome positioning and 5´ transcript length observed in *tpf* and *minu* mutants. TPF-occupied genes that displayed an upstream +1 nucleosome shift of at least 5 pb in both *tpf* and *minu* were ranked by the average shift, and then classified into five equally sized groups. D) Browser screenshots of the four genes studied in Figure 5D showing the 5´extended accumulation of RNA-seq reads upstream the TSS in *tpf* and *minu* mutants in comparison with Col-0. Values indicate normalized read counts.
